# Mitochondrial DNA Mutations Determine Favourable Molecular Responses to Targeted Kinase Inhibitor Therapy and Impair Oxidative Phosphorylation

**DOI:** 10.1101/2025.07.15.664865

**Authors:** Ilaria S Pagani, Vaidehi Krishnan, Ouyang Fengcong John, Chung H Kok, BeiJun Chen, Kian Leong Lee, Cong M Pham, Paul Wang, Elyse Page, Phuong Dang, Verity A Saunders, Jane James, Kelly Lim, Naranie Shanmuganathan, Agnes Yong, Susan Branford, Charles Chuah, David T Yeung, Deborah White, G. Vignir Helgason, Daniel Thomas, Sin Tiong Ong, Timothy P Hughes, David M Ross

## Abstract

Somatic mutations in mitochondrial DNA (mtDNA) are not typically considered key oncogenic drivers of cancer, primarily because of a high synonymous to non-synonymous variant ratio. Here, we surveyed 248 matched diagnosis and remission samples from patients with chronic myeloid leukemia (CML) and found a 75% had mitochondrial mutations with a median number of 2 mutations per patient. mtDNA mutations were predominantly non-synonymous, enriched in the D-loop control region, and likely originated from replication and transcriptional errors. Functionally, mtDNA mutations were associated with reduced oxidative phosphorylation (OXPHOS), as measured by Seahorse analyser. This metabolic vulnerability could be phenocopied by treatment with the complex I inhibitor IACS-10759 in combination with the targeted tyrosine kinase inhibitor (TKI) imatinib, which significantly reduced the colony-forming potential of TKI resistant leukemic stem/progenitor cells (LSPCs). Strikingly, we show that mtDNA mutations were associated with increased sensitivity to imatinib therapy in the clinic. Patients with ≥3 mutations and patients with mutations in the D-loop showed significantly higher cumulative incidence of major molecular response at 24 months (90% vs. 68%, p = 0.004, and 89% vs 68%, p = 0.004 respectively). Single-cell RNA sequencing further revealed enrichment in non-synonymous mtDNA variants in LSPCs from TKI-sensitive patients, while TKI-resistant cells exhibited upregulated gene signatures related to glycerolipid and phospholipid metabolism and mitochondrial biogenesis. Together, our findings demonstrate that mtDNA mutations are key determinants of sensitivity to targeted therapy, rather than oncogenic drivers of leukemogenesis. Mechanistically, non-synonymous mtDNA mutations appear to restrict mitochondrial metabolic plasticity, with widespread implications for precision oncology.

## Introduction

Chronic Myeloid Leukemia (CML) is a myeloproliferative disorder characterized by the t(9;22)(q34;q11) reciprocal translocation, resulting in the *BCR::ABL1* fusion gene on the Philadelphia chromosome.^1^ The BCR::ABL1 oncoprotein is a constitutively active tyrosine kinase that promotes uncontrolled cell proliferation and inhibits apoptosis.^2^ The introduction of Tyrosine Kinase Inhibitors (TKIs) has significantly improved the 5-year survival rate to over 80%.^3^ However, TKI resistance and disease progression to blast crisis remain major contributors to CML-related mortality.^4^ Furthermore, only about 25% of all CML patients can cease their TKI, achieving treatment-free remission, while the remaining patients require lifelong TKI therapy to prevent relapse, leading to organ toxicities, poor drug-adherence, and reduced quality of life.^5,6^

Resistance to TKIs is a complex and multifactorial phenomenon that significantly compromises treatment efficacy in CML.^7^ It can arise through mechanisms such as mutations in the *BCR::ABL1* kinase domain,^8^ acquisition of additional-genetic abnormalities,^9–11^ *BCR::ABL1* overexpression, activation of alternative signalling pathways,^12^ and protective interactions within the bone marrow microenvironment.^13,14^ Metabolic reprogramming has emerged as a key driver of TKI resistance, with leukemic stem and progenitor cells (LSPCs) displaying enhanced oxidative phosphorylation (OXPHOS).^15,16^ These metabolic adaptations confer resistance by enabling CML cells to survive TKI treatment, evade apoptosis, and persist as minimal residual disease, ultimately contributing to relapse and disease progression.^17,18^ Understanding the metabolic dependencies of these cells is crucial for developing novel therapeutic strategies to target resistant LSPCs and potentially achieve a functional cure.

Mitochondria play a central role in this metabolic shift, serving as hubs for OXPHOS-driven ATP production, redox homeostasis, and apoptotic regulation.^19^ The OXPHOS machinery, which is integral to the mitochondrial electron transport chain (ETC), is governed by dual genomic control: while the nuclear genome encodes the majority of OXPHOS proteins, the mitochondrial genome (mtDNA) encodes 13 essential components of Complexes I, III, IV, and V (ATP synthase), alongside 22 transfer RNAs (tRNA) and two ribosomal RNAs (rRNA).^20^ MtDNA is a circular, double-stranded molecule of approximately 16.6 kilobases (kb) which exists in multiple copies per mitochondrion, with each cell containing hundreds to thousands of mtDNA molecules.^21^ The coexistence of identical mtDNA sequences within a cell is referred to as homoplasmy, whereas heteroplasmy describes the presence of both wild-type and mutated mtDNA within the same cell. The threshold level of mtDNA variants can influence cellular function, with high heteroplasmy levels often leading to impaired OXPHOS activity, altered reactive oxygen species (ROS) production, metabolic dysfunction, and increased susceptibility to apoptosis.^22–27^

Although not traditionally regarded as oncogenic drivers, mtDNA mutations are increasingly recognized for their capacity to influence cancer progression and treatment response.^28^ Mutations affecting protein-coding regions of the ETC and the regulatory D-loop region have been reported across diverse malignancies, where they contribute to altered OXPHOS, redox imbalance, and metabolic rewiring.^23,29^ These alterations can enhance cellular adaptability to stress and influence therapeutic response. However, the role of mtDNA mutations in CML remains largely unexplored.

In this study, we present the first comprehensive analysis of mtDNA somatic mutations in paired diagnostic and remission samples from CML patients. We show that somatic mtDNA mutations are prevalent at diagnosis and enriched in protein-coding and regulatory regions critical for mitochondrial function. Importantly, we link these mutations to reduced OXPHOS capacity and increased sensitivity to imatinib, implicating mtDNA integrity as a previously underappreciated determinant of metabolic heterogeneity and therapeutic response in CML. Collectively, we highlight a link between treatment responses and mtDNA mutations in CML, a paradigm for kinase-driven leukaemia. Additionally, our findings position mtDNA mutational profiling as a potential predictive biomarker of TKI response and offer a mechanistic basis for combination strategies aimed at disrupting mitochondrial integrity to overcome drug resistance.

## Results

### Mitochondrial respiration inversely correlates with TKI responses in patient-derived CML LSPCs

To investigate mechanisms underlying variations in TKI response, we profiled mitochondrial function in CD34L LSPCs collected at diagnosis from patients with distinct clinical outcomes. Patients were classified as TKI-sensitive (TKI-S; achieving major molecular response [MMR], *BCR::ABL1* ≤ 0.1% International Scale at 12 months) or TKI-resistant (TKI-Res; failing to achieve MMR at 12 months, progressing to blast crisis, or developing *BCR::ABL1* kinase domain mutations) (**Supplementary Table 1**).

Using the Seahorse XF Analyzer, we measured OXPHOS capacity in CD34+ LSPCs from four TKI-S and five TKI-Res patients at diagnosis, compared to normal CD34+ cells from four G-CSF mobilized healthy donors. Compared with TKI-S, TKI-Res cells exhibited elevated basal oxygen consumption rate (mean 87 vs. 54 pmol/min/10^5 cells, p = 0.036, ordinary one-way ANOVA), maximal respiration potential (mean 213 vs. 115 pmol/min/10^5 cells, p = 0.003, ordinary one-way ANOVA), and spare respiratory capacity (mean 126 vs. 53 pmol/min/10^5 cells, p=0.001, ordinary one-way ANOVA) (**Figure 1A-D**). ATP-linked respiration was also significantly higher in TKI-Res cells compared to TKI-S and normal CD34L cells (mean 78 vs. 51 vs. 6 pmol/min/10L cells, respectively), reflecting enhanced mitochondrial ATP production in resistant LSPCs (**Figure 1E**). These data indicate that TKI-Res LSPCs possess enhanced mitochondrial fitness and metabolic flexibility, a hallmark of increased metabolic adaptability and resilience, in contrast to the relatively lower OXPHOS capacity observed in TKI-S cells.

**Figure 1.**
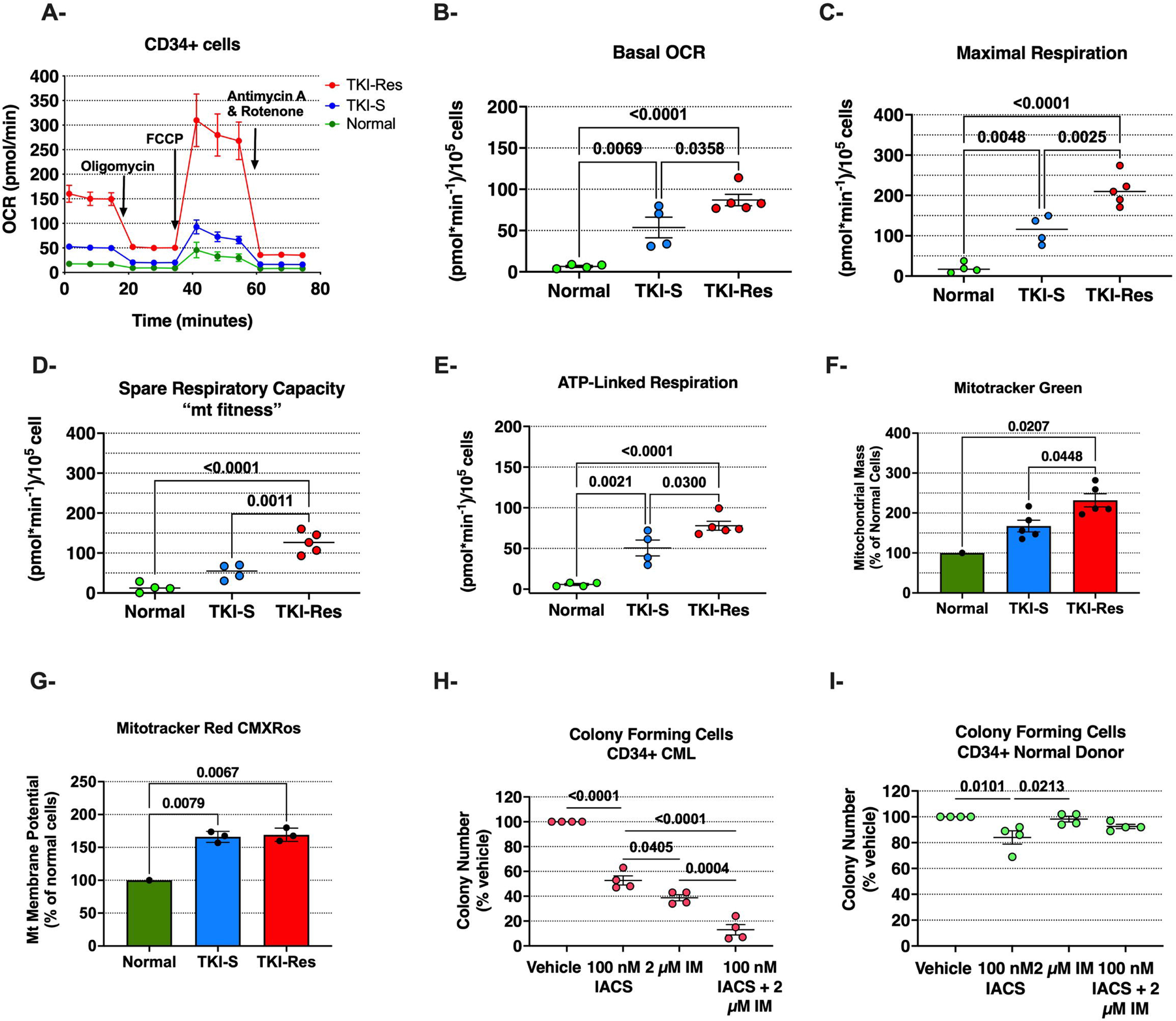
Mitochondrial function in CD34+ cells. **(A)** Representative respirometry output in CD34+ LSPCs (mean with SEM). **(B)** Basal oxygen consumption rate (OCR), **(C)** maximal respiration after injection of Carbonyl cyanide-4 (trifluoromethoxy) phenylhydrazone (FCCP), **(D)** spare respiratory capacity (measured as difference of maximal respiration and basal respiration), and **(E)** ATP-linked respiration (after injection of oligomycin, which inhibits the mitochondrial ATP synthase, or complex V) of four normal HSPCs (green), four TKI-sensitive (blue) and five TKI-Resistant CD34+ LSPCs (red) at diagnosis (ordinary one-way ANOVA, mean with SEM). **(F)** Mitochondrial content was assessed by the median fluorescence intensity (MFI) of Mitotracker Green-labeled CD34+ CML cells (five TKI-sensitive, five TKI-resistant), normalized to that of normal control cells (seven healthy donors) (ordinary one-way ANOVA, mean with SEM); **(G)** Membrane potential was measure by the MFI of MitoTracker Red CMXRos -labeled CD34+ CML cells (three TKI-sensitive and three TKI-resistant), normalized to that of normal control cells (five healthy donors) (ordinary one-way ANOVA, mean with SEM). **(H)** Colony numbers following 3 days of drug treatment of CD34+ LSPCs (four CML patients) (ordinary one-way ANOVA, mean with SEM) **(I)** Colony numbers following 3 days of drug treatment of CD34+ HSPCs (four healthy controls) (ordinary one-way ANOVA, Mean with SEM). *OCR, oxygen consumption rate; Mt, mitochondria*.

Flow cytometry analysis with MitoTracker Green revealed a 1.4-fold increase in mitochondrial content in CD34+ LSPCs from five TKI-Res patients compared to five TKI-S (p = 0.045; ordinary one-way ANOVA; **Figure 1F)**. Mitochondrial membrane potential was significantly higher in both CML populations compared to normal HSPCs (1.6-fold increase, p = 0.008 and p = 0.007, respectively, ordinary one-way ANOVA) but did not differ significantly between the two CML groups (**Figure 1G, and Supplementary Figure 1)**.

Given the enhanced mitochondrial function and metabolic flexibility observed in TKI-Res cells, we tested the efficacy of IACS-010759,^30^ a potent OXPHOS inhibitor of mitochondrial complex I to overcome TKI resistance. Treatment of TKI-Res and TKI-S LSPCs with low-dose (100 nM) IACS-010759 significantly reduced colony-forming potential compared to the vehicle control (p<0.0001, Ordinary one-way ANOVA), with nearly complete eradication of colonies when combined with 2 µM imatinib (p=0.0004, ordinary one-way ANOVA) (**Figure 1H**). While combination therapy with IACS-010759 and imatinib did not affect the number of normal HSPC colonies, IACS-010759 alone induced a modest, albeit statistically significant, reduction in colony formation in normal CD34+ cells (p=0.0101, ordinary one-way ANOVA) (**Figure 1I**).

### Single-cell transcriptomics identifies distinct metabolic reprogramming in imatinib-sensitive and -resistant LSPCs

Single cell RNA-sequencing (scRNA-seq) enables high-resolution analysis of mitochondrial gene expression and mtDNA variants, providing novel insights into cancer-specific metabolic alterations and clonal heterogeneity.^31^ To investigate transcriptional programs associated with mitochondrial function and mtDNA variation in TKI-sensitive and TKI-resistant LSPCs, we re-analysed the publicly available 10X 5’ sc-RNA seq dataset generated by *Krishnan V. et al*.^12^ (**Figure 2A, B**). CML patients in chronic phase (CP) from the Singapore General Hospital (n=17) were categorized into IM-S and two different IM-Res groups based on their response to imatinib, as defined by the European LeukemiaNet criteria.^32^ Diagnostic bone marrow samples were collected prior to TKI treatment. Good responders achieved MMR within 12 months of imatinib treatment and/or deep molecular response (DMR; MR4.5) (CP^MMR^ or Group A). Poor responders were patients who either did not meet molecular (n = 6) or cytogenetic (n = 3) response thresholds on imatinib treatment by 18 months but responded to 2nd/3rd-line TKI (CP^IM-res^ or Group B). Poor responders also included patients who failed all administered lines of TKI therapy and progressed to blast crisis (CP^Fail^ or Group C).

**Figure 2.**
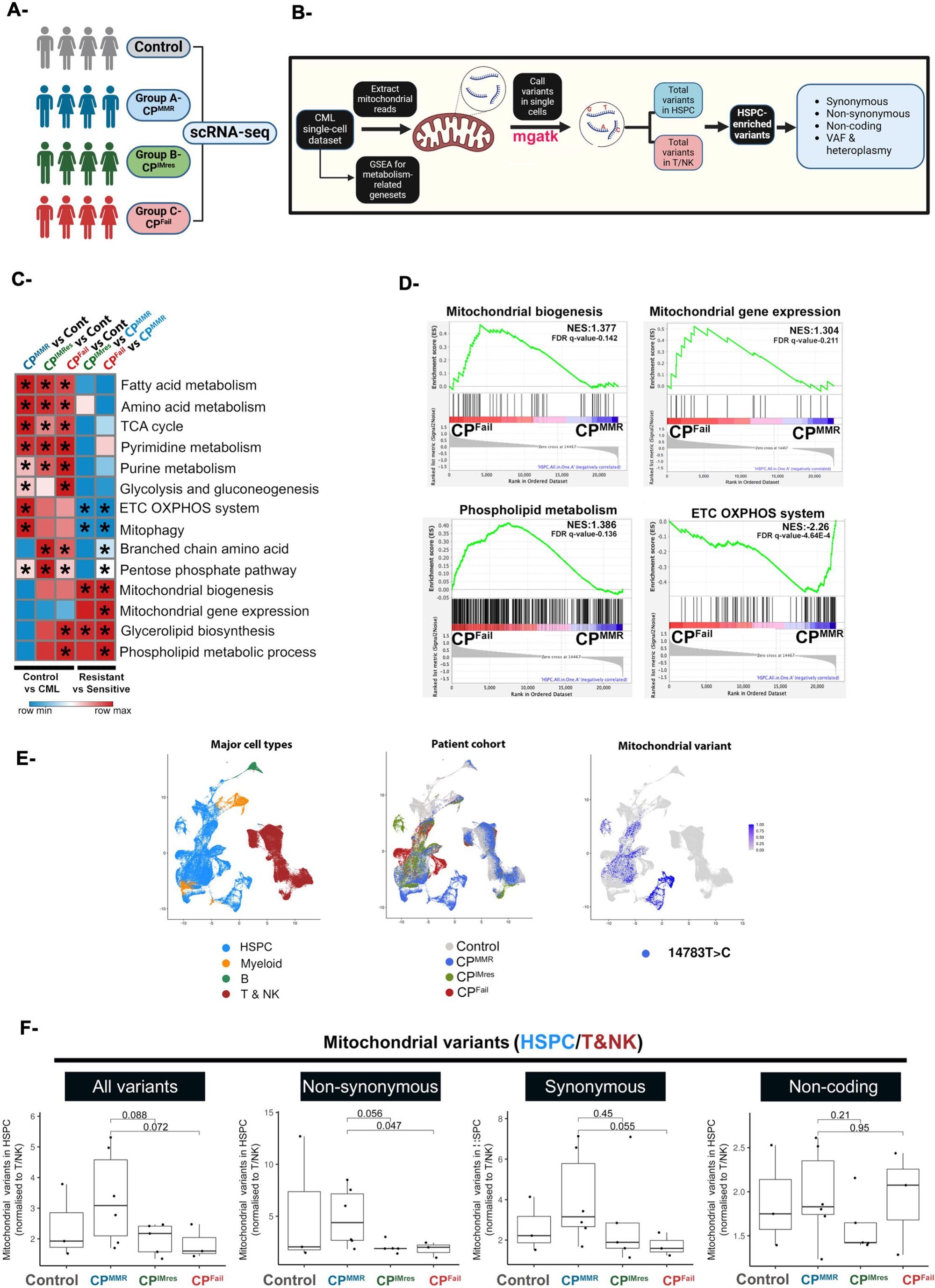
Single-cell transcriptomics reveals distinct metabolic programs and differential mtDNA variant enrichment in imatinib-sensitive and -resistant LSPCs. **(A)** Classification of CML patients (n = 20) from the *Krishnan et al*.^12^ cohort into three groups based on imatinib response: Group A (CP^MMR^; IM-sensitive, optimal response, blue), Group B (CP^IMRes^; sub-optimal response: failure to achieve MMR by 6 months, green), and Group C (CP^Fail^; IM-resistant, treatment failure, red), as per European LeukemiaNet criteria.^32^ Health donors are represented in grey. **(B)** Schematic overview of the analysis pipeline used to extract and annotate mtDNA variants from single-cell 10X 5’ scRNA-seq data. **(C)** Heatmap showing normalized enrichment scores (NES) from gene set enrichment analysis (GSEA) of curated metabolic gene sets in LSPCs (n=20) compared to healthy donor CD34L cells (n=3), and from IM-S IM-Res. Red indicates positive enrichment. **(D)** GSEA comparing LSPCs from TKI-resistant (Groups C) and TKI-sensitive (Group A) patients reveals differential enrichment of mitochondrial and metabolic pathways. upregulation of mitophagy-related gene sets, consistent with mitochondrial impairment. **(E)** UMAP projections of single-cell transcriptomes from bone marrow, coloured by major cell type, and patient group. mtDNA variant calls were mapped onto these clusters to assess distribution across hematopoietic lineages and disease response categories. **(F)** Relative mtDNA variant burden in leukemic stem/progenitor cells (LSPCs), normalized to T and NK cells as internal controls. IM-sensitive patients (Group A) displayed a significantly higher load of non-synonymous mtDNA mutations compared to sub-optimal (Group B, p = 0.056) and resistant patients (Group C, p = 0.047; two-sided t-test).

We performed a comprehensive single-cell transcriptomic analysis to interrogate gene expression programs associated with metabolic reprogramming underlying TKI response phenotypes (**Supplementary Table 2**). Using gene set enrichment analysis (GSEA) against a rigorously curated panel of metabolic and mitochondrial pathways (**Supplementary Table 3**), we compared CD34L LSPCs from CML patients to those from healthy donors. In agreement with previous reports,^15,33^ LSPCs displayed marked enrichment of gene signatures related to fatty acid biosynthesis, amino acid metabolism, and nucleotide synthesis pathways. Additionally, gene sets associated with mitochondrial functions, including the tricarboxylic acid cycle, glycolysis, gluconeogenesis, and branched chain amino acid (BCAA) metabolism were also significantly enriched in LSPCs (**Figure 2C-D**).

We next investigated whether distinct metabolic signatures could differentiate TKI-S and TKI-Res patients. GSEA revealed significant upregulation of gene signatures related to mitochondrial biogenesis and mitochondrial gene expression in TKI-Res samples (**Figure 2C-D**, **Supplementary Figure 2 and Supplementary Table 3**).The latter refers to the coordinated transcription, RNA processing, and translation of mitochondrial-encoded genes, supported by nuclear-encoded regulators essential for mitochondrial function. This transcriptional profile is consistent with enhanced OXPHOS capacity, likely reflecting increased mitochondrial biomass and ATP production. Leading-edge analysis reinforced these findings, highlighting consistent enrichment of genes involved in mitochondrial biogenesis and transcriptional programs, alongside upregulation of phospholipid metabolic processes, glycerolipid biosynthesis, BCAA metabolism, and the pentose phosphate pathway (**Supplementary Figure 2**), indicating a coordinated metabolic adaptation that may contribute to imatinib resistance through preserved mitochondrial function and anabolic flexibility. In contrast, TKI-S samples exhibited downregulation of mitochondrial biogenesis and lipid biosynthetic pathways, alongside increased expression of mitophagy-related gene signatures and ETC OXPHOS system, the latter potentially representing a compensatory mechanism to counteract impaired mitochondrial function.^23^

### Single-cell transcriptomics uncovers enrichment of mtDNA variants in imatinib-sensitive LSPCs

In this study, the median sequencing coverage per cell was 17.7 reads, resulting in a total of 957,796 mtDNA-specific reads mapped across all cells (**Supplementary Figure 3A**). To extract mtDNA variants, we utilized the MGATK (Mitochondrial Genome Analysis Toolkit) pipeline^34^, adapting it for single-cell data processing from the 10X 5’ scRNA-seq platform as described in the methods section. This approach enabled the identification and annotation of mtDNA variants across 54,041 bone marrow cells, facilitating a comprehensive analysis of variant prevalence and heterogeneity among patient groups (**Figure 2E**). To control for background mitochondrial variation and reduce technical artifacts inherent in scRNA-seq, we normalized the number of mtDNA variants detected in LSPCs by calculating their ratio relative to variants found in T and NK cells, which served as reference controls (**Supplementary Figure 3B-E**). Using this normalization, we identified a total of 429 mtDNA variants across 41,290 LSPCs, with a median of 96 variants per patient (**Supplementary Table 4)**.^34^

We observed an enrichment of mtDNA variants in IM-S patients compared to IM-resistant groups (**Figure 2F)**. Notably, non-synonymous sc-mtDNA variants were significantly enriched in LSPCs from IM-S patients compared to those with treatment failure (p = 0.047) with a trend toward enrichment compared to suboptimal responders (p = 0.056).

### mtDNA mutations are highly prevalent in total leukocytes of CML patients at diagnosis

Building on our observation that mtDNA variants are enriched in LSPCs from imatinib-sensitive patients, we next investigated the broader landscape of mitochondrial genome alterations in CML. Somatic mtDNA mutations have been increasingly recognized as contributors to cancer biology, with potential to modulate oxidative phosphorylation, redox homeostasis, and apoptotic signaling.^23,28,29^ However, their role in CML remains largely unexplored. In fact, whether mtDNA mutations may underpin the observed heterogeneity in mitochondrial respiration and treatment responses in CML cells is unknown.

To address this knowledge gap, we performed a comprehensive analysis of mtDNA somatic point mutations in peripheral blood total white cells (TWCs) from CML patients at diagnosis, using matched samples (either non-hematopoietic cells or TWCs in MMR collected at 12 months of TKI therapy) as germline controls.^35^ This approach enabled us to identify mtDNA somatic point mutations by filtering out pre-existing germline variants. We identified 244 somatic mutations in 93/124 (75%) of CML patients at diagnosis (**Supplementary Table 5**). The median number of mutations per patient was 2, ranging from 1 to 21. Notably, four patients carried ≥ 10 mtDNA mutations (**Figure 3A**). Most of the substitutions were heteroplasmic with median variant allele frequency, VAF 9.6%. Only 29 (12%) were homoplasmic substitutions (VAF ≥ 98%) and only one additional variant had a VAF exceeding 50% (**Figure 3B**). The predominance of heteroplasmic mutations with lower VAF suggests that these mutations are sub-clonal and not under strong positive selection prior to TKI treatment.^36^ In contrast, the presence of mutations with higher VAF could reflect either an origin early in the evolution of CML or a positively selected clonal expansion.^21^

**Figure 3.**
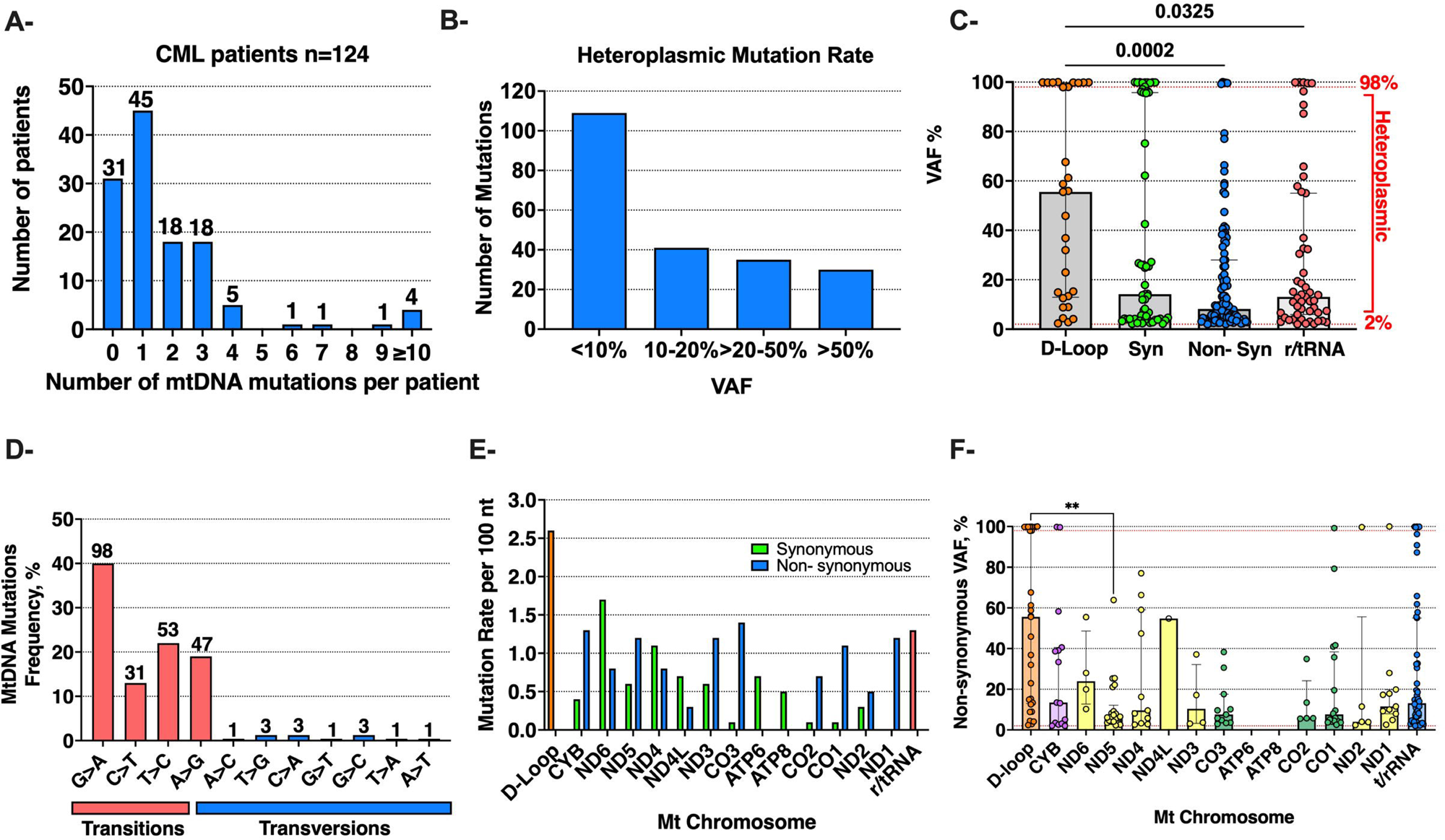
MtDNA mutations in CML patients at diagnosis. **(A)** Number of mtDNA somatic mutations per patient identified in 124 CML patients at diagnosis. **(B)** Variant allele frequency (VAF) distribution of heteroplasmic somatic mtDNA mutations. **(C)** VAF of mutations across different regions of the mitochondrial genome. Mutations with VAF between 2% and 98% are considered heteroplasmic, while mutations with VAF greater than 98% are considered homoplasmic. (Kruskal-Wallis test; median with interquartile range). **(D)** Frequency of mutation types across the mtDNA genome. The majority of mutations were transitions (red), with G:C>A:T mutations comprising 40% and T:A>C:G mutations 22%. A smaller proportion of mutations were transversions (blue). **(E)** Mutation rate per 100 nucleotides in different mtDNA regions. The mutation rate per 100 nucleotides was calculated by dividing the total number of mutations identified in each mtDNA region by the total number of nucleotides in that region. The plot shows the mutation rates for various mtDNA regions, including the D-loop (orange), structural RNAs (rRNA and tRNA, red) and coding regions, highlighting the relative mutation frequencies across these areas. Mutations in the coding region were further subdivided in synonymous (green) and non-synonymous (blue). **(F)** Distribution of VAFs of non-synonymous mutations in mitochondrial genes (Kruskal-Wallis test; median with interquartile range).

The distribution of somatic point mutations across the mitochondrial genome is shown in **Figure 3C**. Non-coding mutations in the D-loop control region (29 mutations in 9 patients) exhibited the highest VAF, with a median of 56%, reflecting a high prevalence of homoplasmic mutations in this region. The majority of the substitutions were transitions, with G:C>A:T (40%) and T:A>C:G (22%) being the most frequent mutation types (**Figure 3D)**. This pattern is most consistent with errors introduced by DNA polymerase gamma (Polγ) during mtDNA replication.^28,37^

In coding regions, we detected 107 non-synonymous and 57 synonymous somatic mutations. Notably, *MT-ND1* (*NADH dehydrogenase 1*, Complex I) exhibited 10 non-synonymous and no synonymous mutations (p = 0.016, Fisher’s exact test; Bonferroni-adjusted p = 0.202), suggesting strong positive selection in this region (**Figures 3E, F**). Similarly, *MT-CO1* (*cytochrome c oxidase subunit I*, Complex IV) showed a skewed ratio of non-synonymous to synonymous mutations (17 vs. 3; p = 0.02, Bonferroni-adjusted p = 0.264), possibly indicating selective pressure favouring protein-altering changes. In contrast, all mutations identified in *ATP6* and *ATP8* (*ATP synthase membrane subunits 6 and 8*, Complex V) were synonymous, suggesting negative selection against amino acid–altering variants in ATP synthase.^23^ Although the number of mutations per gene is modest and subgroup sample sizes are limited, these trends suggest non-neutral mutational dynamics and warrant validation in larger, independent cohorts.

### High variant allele frequency and homoplasmic mtDNA mutations are enriched in imatinib-sensitive CML patients

To explore whether mtDNA load affects treatment responses in CML, we focused on a subset of patients treated with imatinib stratified as IM-sensitive (IM-S; n = 41) or IM-resistant (IM-Res; n = 39) (**Supplementary Table 1)**.

MtDNA somatic point mutations were detected in 31 (76%) IM-S and 29 (74%) IM-Res patients, indicating a similar proportion of affected individuals in both groups. However, IM-S harboured a total of 139 mutations (median 3 mutations/patient, range 1–21) compared to 39 in the IM-Res group (median 1 mutation/patient, range 1-3; p = 0.062, Mann-Whitney test) (**Figure 4A**). Notably, mutations detected in IM-S had significantly higher VAF than those in IM-Res (median 21% vs. 7%, p < 0.0001, Mann-Whitney test) (**Figure 4B, C**). All the 29 homoplasmic mutations were identified in IM-S patients, while IM-Res exhibited exclusively heteroplasmic mutations, the majority of which had VAF < 20% (**Figure 4D**).

**Figure 4.**
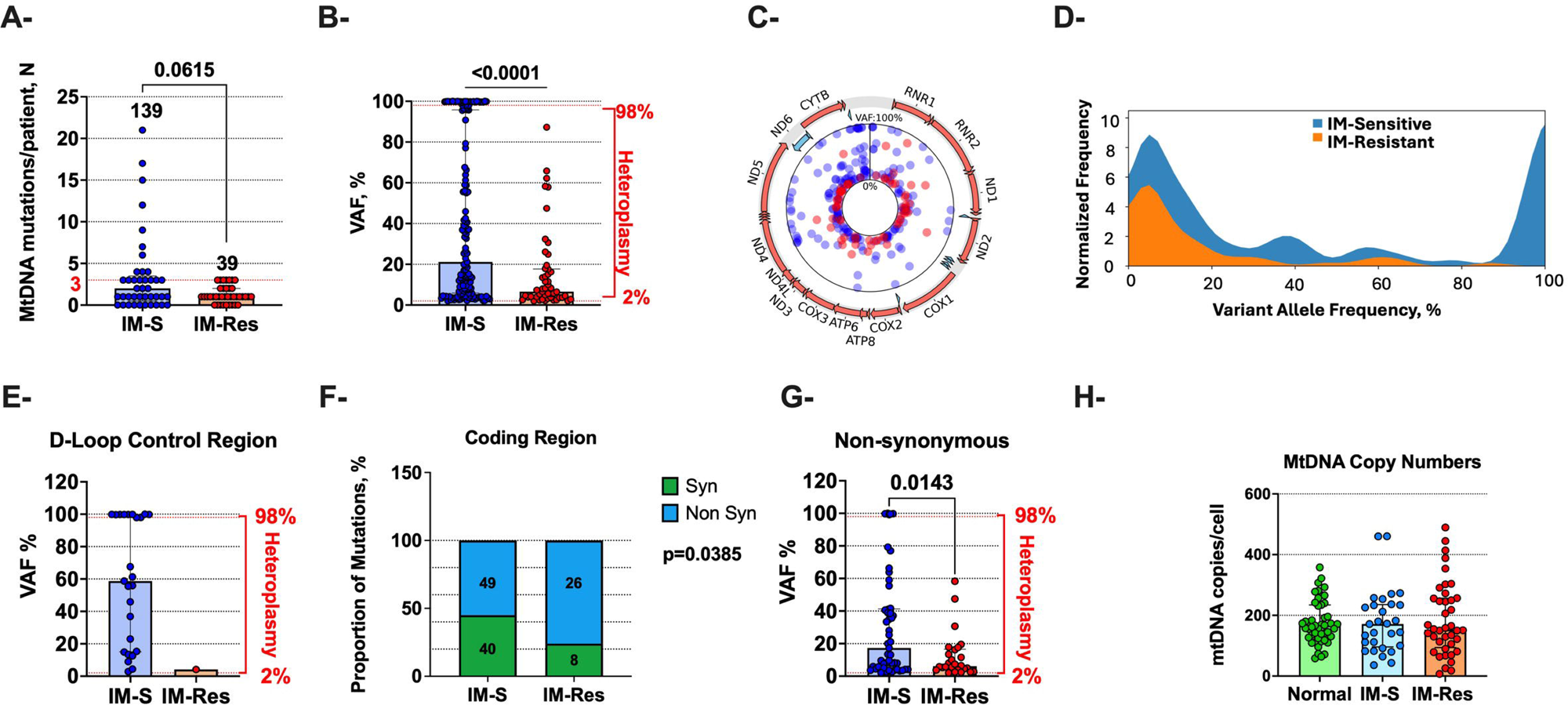
Association between mtDNA mutations and response to imatinib (IM). **(A)** Mutational burden in IM-sensitive (IM-S, blue) and IM-resistant (IM-Res, red) patients (Mann-Whitney U test, median with interquartile range). **(B)** VAF of mtDNA mutations in IM-S and IM-Res patients (Mann-Whitney test, median with interquartile range). **(C)** Circos plot representing the landscape of mtDNA mutations in IM-S (blue) and IM-Res (red) patients. The dots represent the VAF distribution from 2% (centre) to 100%, illustrating the prevalence and clonal expansion of mutations in each group. **(D)** Distribution of VAF for mtDNA mutations in IM-S (blue) and IM-Res (orange) patients. IM-S patients exhibited a higher proportion of mutations with VAF > 10%, including mutations with VAF > 50%. In contrast, IM-Res patients had a greater proportion of low-VAF mutations (<10%). **(E)** VAF of mtDNA mutations in the D-Loop control region between IM-S and IM-Res patients (Mann-Whitney U test, median with interquartile range). **(F)** Proportion of synonymous (green) and non-synonymous (blue) mutations in IM-S and IM-Res patients (Fisher’s Exact Test). **(G)** VAF of non-synonymous mutations in IM-S and IM-Res patients (Mann-Whitney U test, median with interquartile range). **(H)** Boxplot of mtDNA copy number per cell in normal (n=47, green), IM-S (n=29, blue) and IM-Res (n=40, red). Data were calculated as the ratio of mtDNA (MT-ND1 and MT-CYB)/nDNA (GUSB) by PCR. (Kruskal-Wallis test, mean with SEM).

To further explore these findings, we analysed the distribution of mutations across distinct mitochondrial regions, including the D-loop control region, coding sequences, and structural RNA (r/tRNA). In IM-S patients, a significantly higher proportion of mutations were observed in the D-loop region compared to IM-Res patients (25/139 vs. 1/49, p < 0.0001, Fisher’s exact test comparing the distribution of D-loop vs. non–D-loop mutations between response groups; (**Fig. 4E**). This indicates that regulatory elements in the mitochondrial genome may represent a mutational hotspot associated with treatment response, consistent with previous observations of the D-loop as a mutation-prone region in cancer-associated mitochondrial genomes.^23^ In contrast, no differences were observed in the frequency or VAF of mutations within structural RNA (r/tRNA) regions, implying a lack of selective pressure in this domain (**Supplementary Fig. 4A**).

Furthermore, IM-Res patients exhibited a skewed distribution favouring non-synonymous over synonymous mutations (p = 0.039, Fisher’s exact test; **Figure 4F**). However, non-synonymous mutations in IM-S patients occurred at significantly higher VAF than those in IM-Res (median 17% vs. 6%; p = 0.014, Mann–Whitney test; **Figure 4G**), indicating that a higher clonal burden of protein-altering mutations may be associated with imatinib sensitivity. No significant difference was observed in the VAF of synonymous mutations between the groups (**Supplementary Figure 4B**).

There was no significant difference in the predicted pathogenicity of non-synonymous mutations between IM-S and IM-Res, with both groups exhibiting similar distributions of high PolyPhen and deleterious SIFT scores (**Supplementary Figures 4C and D**). IM-Res patients harboured a greater number of non-synonymous mutations, but these may remain largely sub clonal. In contrast, IM-S patients exhibited enrichment of non-synonymous mutations at higher VAFs, indicating clonal dominance and potentially greater mitochondrial dysfunction. Therefore, these findings support a model in which mtDNA-driven mitochondrial impairment may contribute to increased therapy response rate in CML patients.

To explore whether mtDNA mutations may underlie the metabolic heterogeneity observed in **Figure 1**, we analysed matched TWCs samples from the same patients profiled by Seahorse assays (**Figure 1A–D; Supplementary Table 6**). TKI-sensitive patients, who exhibited lower OXPHOS capacity, had a higher mtDNA mutation burden (median 2 mutations per patient, range 1–12) with significantly higher VAFs (median 56%, range 2.5 – 97.1%) compared to TKI-resistant patients, who showed elevated OXPHOS (median 1 mutation, range 1–2; median VAF 5.5%, range 4.2–8.2%; p = 0.004, Mann–Whitney test). These findings mirror trends in the broader cohort and support a link between high-VAF mtDNA mutations and reduced mitochondrial respiration in TKI-sensitive CML cells.

We next investigated whether the observed mutational differences between IM-S and IM-Res were associated with changes in mtDNA abundance in the bulk TWCs (**Figure 4H**). No significant differences were observed in mtDNA/nDNA ratios as determined by Q-PCR (median mtDNA copies/cell 171 vs 152 in IM-S and IM-Res; Kruskal-Wallis test), likely reflecting the cellular heterogeneity of bulk leukocyte populations.

### MtDNA mutations in CML patients are associated with better response to imatinib

We finally assessed the impact of mtDNA mutations on imatinib molecular response. A cutoff of ≥3 mutations was chosen based on the observation that IM-Res patients harboured no more than 3 mutations. Patients with ≥3 mutations showed significantly higher cumulative incidence of MMR and MR4.5 at 24 months (90% vs. 68%, p=0.004; and 63% vs. 37%, p = 0.011, respectively; **Figure 5A, B**).

**Figure 5.**
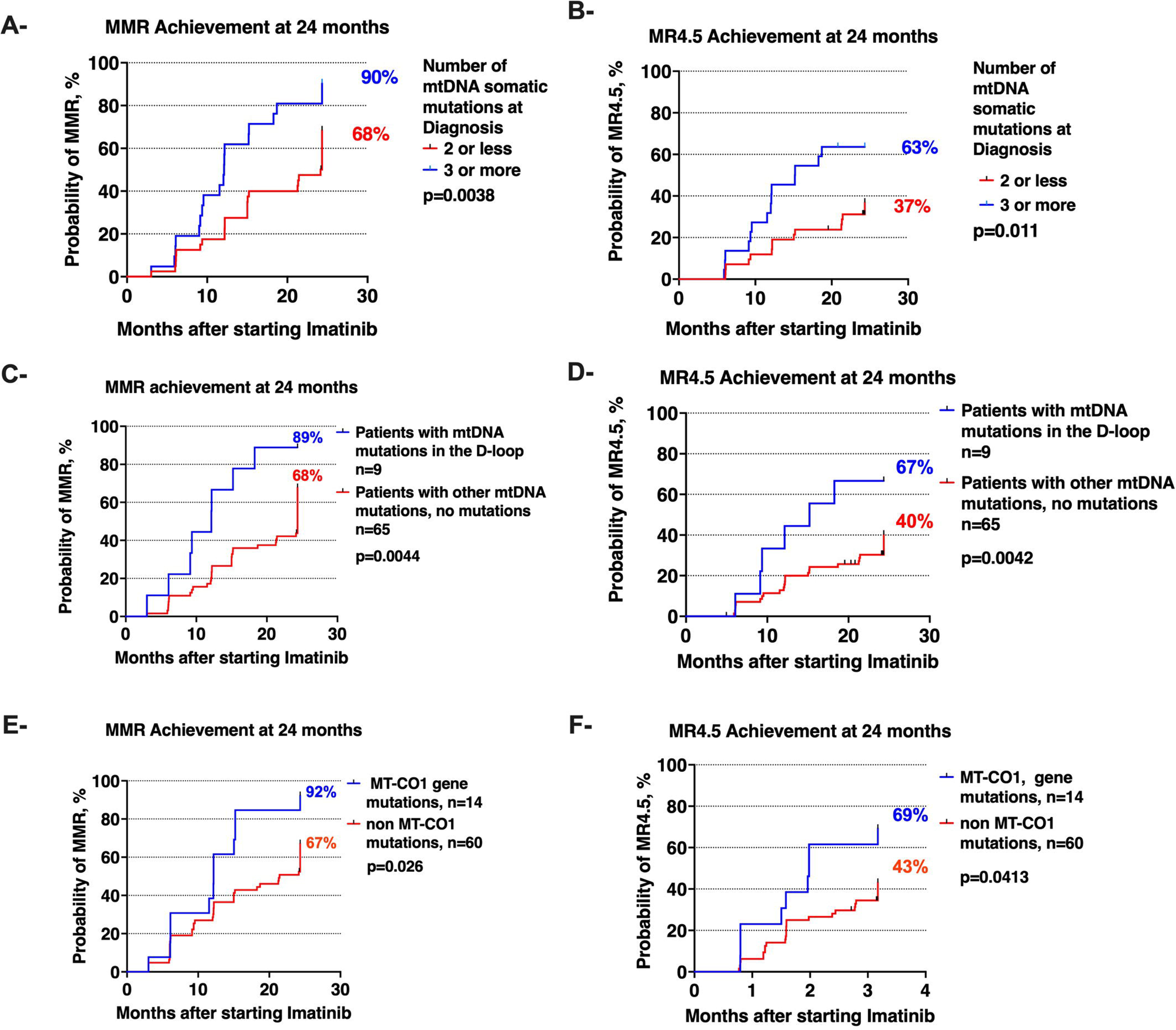
Impact of mtDNA mutations on imatinib response. Kaplan-Meier analysis of cumulative incidence of MMR and MR4.5 at 24 months in relation to mtDNA mutations. **(A)** MMR related to mutational burden, ≥3 or <3 mutations (90% vs 68%; hazard ratio log rank [HR] 2.23; 95% CI, 1.34 – 4.39) and **(B)** MR4.5 (63% vs 37%; hazard ratio log rank [HR] 2.47; 95% CI, 1.09 – 5.59). **(C)** MMR related to D-Loop mutations vs non-D-loop (89% vs 68%; hazard ratio log rank [HR] 2.74; 95% CI, 0.89 – 8.42) and **(D)** MR4.5 (40% vs 67%; hazard ratio log rank [HR] 2.38; 95% CI, 2.71 – 8.09). **(E)** MMR related to MT-CO1 mutations vs non-MT-CO1 mutations (92% vs 67%; hazard ratio log rank [HR] 2.04; 95% CI, 0.91 – 4.61) and **(F)** MR4.5 (43% vs 69%; hazard ratio log rank [HR] 2.13; 95% CI, 0.82 – 5.54).

We further analysed the most frequently mutated loci. Mutations in the D-loop (**Figure 5C, D**) and *MT-CO1* (**Figure 5E, F**) had a higher cumulative incidence of MMR (89% vs. 68%, p = 0.004 for D-loop; 92% vs. 67%, p = 0.026 for *MT-CO1*) and MR4.5 at 24 months (67% vs. 40%, p = 0.004 for D-loop; 69% vs. 43%, p = 0.041 for *MT-CO1*).

We also examined overall survival (OS), failure-free survival (FFS), and progression-free survival (PFS) in relation to mtDNA mutations. No significant difference in OS was observed, likely due to the low incidence (only two deaths, both in the IM-Res group). Similarly, no differences in PFS were detected. However, the number of mtDNA mutations showed a trend toward improved FFS, with 95% vs. 75% at 24 months (p = 0.059; **Supplementary Figure 5**), suggesting a potential association of higher mtDNA mutational burden with long-term disease control.

## Discussion

Advances in next-generation and single-cell sequencing have enabled high-resolution analysis of mtDNA alterations, revealing previously underappreciated mitochondrial heterogeneity.^34^ MtDNA mutations are increasingly recognized for their role in cancer metabolism,^20,23,28^ but their relevance in CML has remained largely unexplored. To our knowledge, this is the first study to comprehensively profile mtDNA mutations in CML and link them to metabolic reprogramming and therapy response. While the Warburg effect highlights a shift toward glycolysis in cancer cells, accumulating evidence suggests that therapy-resistant cells often depend on enhanced OXPHOS to meet the energetic and biosynthetic demands of survival.^15,17^ A recent study by *Li-Harms et al.* provided compelling evidence that mtDNA mutation burden can shape metabolic plasticity during leukemogenesis.^38^ Using Polg^D257A mutator mice, they showed that hematopoietic progenitors with a moderate mtDNA mutation load retained the ability to adapt metabolically and support acute leukemic transformation under NMyc stress, whereas those with a high mutation burden exhibited impaired mitochondrial metabolism, reduced metabolic flexibility, and failed to sustain leukemogenesis. These findings are concordant with our observation that mtDNA mutational profiles in CML are associated with divergent metabolic programs and therapy sensitivity.

Our data reveal a substantial burden of somatic mtDNA mutations in newly diagnosed CML patients. These mutations were predominantly heteroplasmic and enriched in functionally critical regions, including the D-loop and genes encoding subunits of respiratory complexes I and IV. Notably, patients who achieved better responses to imatinib harboured mutations at higher VAFs. Homoplasmic mutations could be functionally neutral, but acquired early in the ontogeny of CML, or they could confer selective pressure leading to outgrowth of the clone.^28^ The lack of distinct mutational hotspots in the coding regions of the mtDNA may suggest that the former explanation is more likely. Nevertheless, the difference in sensitivity to imatinib supports the concept of a "heteroplasmic threshold",^21^ where the proportion of mutant mtDNA within the mitochondrial population must exceed a critical level to disrupt OXPHOS efficiency and cellular bioenergetics, leading to mitochondrial dysfunction, and increased cellular stress.

The predominance of G:C>A:T transitions implicates a replication-associated mutational process, rather than oxidative damage.^28^ This signature aligns with the limited proofreading capacity of the Polγ, which is prone to misincorporation errors during replication, and is consistent with the mitochondrial mutational profile reported in other cancers.^20,28,39^ Additionally, the strand-asynchronous model of mtDNA replication may expose the heavy strand to spontaneous deamination,^40^ contributing to the observed transition bias and overall mutational burden.

Our single-cell RNA sequencing data further reveal a distinct mitochondrial transcriptional profile in TKI-sensitive cells, characterized by upregulation of genes involved in the ETC OXPHOS system and mitophagy, alongside suppression of genes controlling mitochondrial biogenesis. The upregulation of OXPHOS genes, in the context of impaired biogenesis, likely represents a compensatory attempt to preserve mitochondrial function.^23^ Similarly, upregulated mitophagy could be a response to accumulated mtDNA damage. This pattern suggests an uncoupling of mitochondrial turnover and renewal, whereby dysfunctional mitochondria may be selectively removed, yet insufficient biogenesis prevents adequate replacement.^41^ The resulting imbalance may lead to a metabolically compromised mitochondrial pool, harbouring deleterious mutations that impair respiratory chain function and lower the apoptotic threshold, rendering these cells more susceptible to TKI-induced cell death.^41^

In contrast, TKI-resistant cells displayed a significantly lower mtDNA mutational burden and lower VAFs, suggesting that the preservation of mitochondrial genomic integrity may contribute to survival of LSPCs in the face of imatinib treatment. This aligns with findings in a Li-Fraumeni syndrome mouse model, where excessive mtDNA mutations, induced by a proofreading-deficient Polγ, impaired tumorigenesis, underscoring the importance of maintaining mitochondrial genomic integrity for leukemic potential.^42^ Transcriptomic analysis and functional assays further revealed that TKI-resistant LSPCs at diagnosis had upregulated genes involved in mitochondrial biogenesis and related to mtDNA transcription, RNA processing, and translation, reflecting an adaptive response to mitochondrial stress.^15^ Additionally, TKI-resistant LSPCs exhibited upregulation of anaplerotic metabolic pathways, including the pentose phosphate pathway and BCAA catabolism. These metabolic adaptations support OXPHOS by supplying NADPH for redox homeostasis and replenishing intermediates of the tricarboxylic acid cycle, thereby sustaining energy production.^43^ Furthermore, enhanced glycerolipid biosynthesis and phospholipid metabolism were observed, which likely support mitochondrial membrane integrity and function. Collectively, these findings delineate a program of metabolic reprogramming in TKI-resistant LSPCs, characterized by enhanced mitochondrial efficiency, bioenergetic plasticity, and redox resilience, all of which contribute to their survival and adaptation under therapeutic pressure.

These findings align with evidence from other malignancies, where mtDNA mutations have been associated with altered cellular metabolism; however, findings regarding treatment outcomes remain inconsistent and context-dependent, with limited functional validation.^20^ For example, in prostate cancer non-synonymous mtDNA mutations in complex I have been associated with adverse outcome, OXPHOS remodelling and altered expression of metabolic enzymes involved in pyruvate and succinate metabolism.^44^ Cybrid models using patient-derived renal carcinoma cells have demonstrated that mutations in *MT-CYB* induce a metabolic shift toward reductive carboxylation of glutamine to support lipid synthesis and proliferation of cancer cells.^25^ In contrast, cybrid studies in melanoma reveal that pathogenic mtDNA mutations impair mitochondrial function and reduce metastatic potential by limiting tumour cell invasion and dissemination.^45^ Other studies report that mutations in the mtDNA D-loop region correlate with improved treatment outcomes in prostate cancer and oral squamous cell carcinoma.^46^ Larger-scale studies integrating clinical outcomes with detailed functional assessments of mitochondrial alterations and function are needed to clarify these relationships and guide therapeutic strategies.

In CML, metabolic reprogramming is further influenced by TKI therapy. Recent studies reveal that TKI treatment induces dynamic changes in energy metabolism of leukemic stem cells, characterized by an initial suppression of OXPHOS, glycolysis, and fatty acid oxidation pathways, which are subsequently restored during prolonged treatment.^17,43^ This metabolic plasticity enables leukemia stem cells to adapt and persist despite TKI therapy, underscoring the critical role of mitochondrial integrity and metabolism in therapeutic resistance.

Targeting mitochondrial function in combination with TKI therapy represents a compelling approach to overcoming resistance in CML, and potentially in other cancers that show response to TKIs. In our preclinical model, the mitochondrial complex I inhibitor IACS-010759 effectively sensitized resistant CML cells to imatinib, highlighting the therapeutic potential of disrupting mitochondrial metabolism. However, translating this approach into clinical practice remains challenging. The Phase I/II trial of IACS-010759 (NCT02735235) was terminated prematurely due to multi-system, dose-limiting toxicities, underscoring the narrow therapeutic window of systemic inhibition of mitochondrial function.^47^ Future strategies may focus on selective delivery systems, alternative mitochondrial targets with improved safety profiles, or transient inhibition approaches to limit off-target toxicity. To address this, a recent drug-repurposing screen identified lomerizine, a calcium channel blocker, as a selective mitochondrial metabolism inhibitor in CML, with reduced systemic toxicity.^48^ Together, these findings support further exploration of mitochondrial vulnerabilities in TKI-resistant CML.

In conclusion, our findings uncover a previously underappreciated role for mtDNA mutations as determinants of sensitivity to targeted therapy, independent of their function as classical oncogenic drivers. By linking non-synonymous mtDNA mutations to impaired mitochondrial plasticity and increased treatment susceptibility, this study reshapes our understanding of how mitochondrial genetics influence therapeutic outcomes. While our focus is CML, these insights may extend to other malignancies treated with targeted agents, where mitochondrial metabolism plays a pivotal role in cell survival and resistance. Our results pave the way for exploiting mitochondrial vulnerabilities in precision oncology, and for developing metabolism-informed strategies to overcome therapy resistance.

## Methods

### Patient characteristics

We retrospectively studied 124 chronic-phase CML patients at diagnosis (pre-TKI) and after 12 months of therapy with either imatinib (nL=L80) or nilotinib (nL=L44) (**Supplementary Table 1**). Patients were enrolled in the following clinical trials: Australian Leukaemia and Lymphoma Group CML6 (TIDEL I; ACTRN12607000614493, *n*L=L13), ALLG CML8 (TIDEL II; ACTRN12607000325404, *n*L=L67), ALLG CML11 (PINNACLE, ACTRN12612000851864, *n*L=L22), and ENESTxtnd (NCT01254188, *n*L=L22).

Imatinib-treated patients included 41 classified as TKI-sensitive and 39 as TKI-resistant (TIDEL I: 6 sensitive, 7 resistant; TIDEL II: 35 sensitive, 32 resistant). Among nilotinib-treated patients, 30 were TKI-sensitive and 14 TKI-resistant (PINNACLE: 17 sensitive, 5 resistant; ENESTxtnd: 13 sensitive, 9 resistant).

### Primary samples

Total leukocytes or mononuclear cells were isolated from peripheral blood of CML patients at diagnosis and after 12 months of TKI therapy,^35^ with informed consent obtained in accordance with the Declaration of Helsinki and approval of Central Adelaide Local Network (HREC/14/RAH/316). Normal samples were obtained from cryopreserved peripheral blood bags of healthy G-CSF-mobilized donors. Matched non-hematopoietic cells including hair follicles and mesenchymal stem cells, were available for a subset of patients.

CD34+ cells were isolated from peripheral blood CML and G-CSF-mobilized mononuclear cells using the CD34 MicroBead Kit (Miltenyi Biotec, Germany) and manual MACS Cell Separation (Miltenyi Biotec, Germany). Following MACS separation, CD34+ cells were cryopreserved for further analysis.

### Cell culture

Primary samples were cultured at 37°C and 5% CO2 in serum-free media (SFM) consisting of Iscove’s modified Dulbecco medium (Sigma-Aldrich, Missouri, United Stated) supplemented with BIT 9500 Serum Substitute (bovine serum albumin, BSA/insulin/transferrin; STEMCELL Technologies, Vancouver, Canada), 2 mM L-Glutamine (Sigma-Aldrich, St. Louis, Missouri, United States), 1 mM streptomycin/penicillin, 0.1 mM 2-mercaptoethanol, 0.02 mg/mL human Low Density Lipoprotein (STEMCELL Technologies, Vancouver, Canada). The SFM was then filtered and supplemented with a physiological growth factor cocktail (STEMCELL Technologies, Vancouver, Canada) consisting of 0.05 ng/mL human recombinant LIF, 1 ng/mL G-CSF and 0.2 ng/mL SCF/GM-CSF/MIP-α/IL-6.

### Drug Treatment

CD34+ cells were incubated with 2 µM Imatinib (kindly donated by Novartis, Basel, Switzerland) and 100 nM IACS-010759 (Selleck Chemicals LLC, Texas, United States), either individually or in combination. IACS-010759 was reconstituted in DMSO prior to use.

### Oxygen consumption rate (OCR) measurement

OCR was measured using the Seahorse XF96 Analyzer (Agilent Technologies, California, United States) with the Seahorse XF Cell Mito Stress Test Kit (Agilent, California, United States). Primary cells were cultured overnight in SFM with physiological growth factors and then resuspended in warm Seahorse XF RPMI medium (pH 7.4), supplemented with 10 mM glucose, 1 mM pyruvate, and 2 mM glutamine (all from Agilent Technologies, California, United States). Eight replicates were performed per sample.

Cells (1.2 × 10L per well) were seeded on Cell-Tak-coated plates and centrifuged at 800 rpm for 3 minutes, followed by a 1-hour incubation at 37°C in a non-COL incubator. OCR was recorded at baseline and after sequential injections of oligomycin (1.5 μM), FCCP (2 μM), and rotenone plus antimycin A (0.5 μM each). Measurements were normalized to cell number. Additional details are provided in **Supplementary Methods**.

### Mitochondrial content and membrane potential

Mitochondrial content and membrane potential were assessed using MitoTracker Green FM and MitoTracker Red CMXRos (Invitrogen Carlsbad, California) fluorescent probes, following the manufacturers’ instructions. Additional details are provided in **Supplementary Methods**.

### Colony Forming Cell (CFC) assays

Primary cells were plated in SFM supplemented with PGF cocktail in the presence of indicated drugs. After 3 days, cells from each condition were transferred in methylcellulose-based medium (MethoCult H4034 Optimum, STEMCELL Technologies) in duplicate and colonies were manually counted after 12-14 days.

### Next-generation sequencing of the entire mitochondrial genome

MtDNA mutations were identified as previously described.^35^ Briefly, genomic DNA was extracted using a phenol/chloroform method. The full 16.9 kb mitochondrial genome was amplified by long-range PCR using overlapping primer sets, and pooled amplicons were subjected to library preparation using the NexteraXT kit (Illumina, San Diego, CA)) and sequenced on an Illumina MiSeq platform. Detailed PCR conditions, primer sequences, and purification steps are provided in the **Supplementary Methods**.

### Calling of mtDNA variants and somatic mutations

Paired-end mitochondrial reads were aligned to the Revised Cambridge Reference Sequence (rCRS, NC_012920), which includes a 3107N deletion to preserve historical position numbering. Alignment was performed using bwa-mem2 v2.2.1, and duplicate reads were marked using PicardTools v3.2.0.

Variant calling was conducted with LoFreq v2.1.5 in somatic mode, using a matched sample taken after 12 months of TKI therapy, or either hair follicle or mesenchymal stem cell from the same patient, as germline control. Functional annotation of variants was performed using the Ensembl Variant Effect Predictor (VEP) web interface, including predictions from SIFT and PolyPhen-2.

We previously empirically identified a cut off of 2% to discriminate between sequencing errors (false positive) and variants at low Variant Allele Frequency (VAF).^35^ All variants with VAF < 2% were removed; variants with VAF 2–98% were considered as heteroplasmic, variants with VAF ≥98% were considered as homoplasmic, and at least five supporting reads per pair-end were required. Several errors were removed from the final list of variants. They included the false positives C3106A, AC3105A, N3107C generated by misalignment due to the 3107N in the rCRS; the homopolymer tract in the region 302–315 generating the recurrent T310C, T310TC misalignments; the T333C and T344C recurrent false positives. Single Nucleotide Polymorphisms (SNPs) were also excluded.

### MtDNA copy number quantification

Relative mtDNA copy number was quantified by real-time quantitative PCR (qPCR) using two mitochondrial gene targets, MT-ND1 (Hs02596873-s1, FAM-MGB) and MT-CYB (Hs02596867-s1, FAM-MGB), normalized to the nuclear reference gene GUSB.^49^ Reactions were performed in triplicate using TaqMan assays on a QuantStudio™ 7 Flex system (Life Technologies, California, USA). The relative copy number per diploid genome was calculated using the ΔCt method, averaging results from both mitochondrial targets. Detailed primers, probes, reaction conditions, and calculation formulas are provided in the **Supplementary Methods**.

### Calling mtDNA variants from scRNA-seq data

To identify variants in the mitochondrial genome at the single-cell level, we employed the Mitochondrial Genome Analysis Toolkit (mgatk).^34^ The mgatk pipeline was executed on our previously published single-cell RNA sequencing (scRNA-seq) dataset^12^ using the following parameters: the ’tenx’ command was used to indicate input 10x Genomics single-cell data. The input BAM file was specified using the ’-i’ flag, followed by the path to the ’possorted_genome_bam.bam’ file for each sample. The output label and directory were set using the ’-n’ and ’-o’ flags, respectively. For accurate cell and molecule identification, we set the unique molecular identifier (UMI) tag to ’UB’ using the ’-ub’ flag, and the barcode tag to ’CB’ using the ’-bt’ flag. The path to the barcodes file was provided using the ’-b’ flag, pointing to the ’barcodes.tsv’ file in each sample’s directory, corresponding to the cell barcodes of cells passing QC and present in the final dataset. Finally, we specified the reference genome as GRCh38 using the ’-g’ flag. The output signac[c] rds object from running the mgatk pipeline was then used for downstream analysis using the signac package (v1.14).

Using the signac package, the ‘IdentifyVariants’ function was used to determine each variant’s coverage and low confidence variants are filtered, keeping variants with n_cells_conf_detected >= 2. Allele frequencies at the single-cell level are calculated using the ‘AlleleFreq’ function. We applied a single-cell allele frequency cutoff of > 0.1 to determine if a cell contained the variant of interest. The %VAF is then determined as the percentage of cells containing the variant of interest within a specific cell type (e.g. HSC, MKP, ERP etc) of a particular patient. Next, we compared the number of variants in hematopoietic stem and progenitor cells (HSPCs) as compared to T&NK cells. As the sequencing depth varies and can affect the number of variants detected, we normalised the number of variants in HSPC against that in T&NK cells by computing the HSPC-to-T&NK variant ratio. Furthermore, for each patient, variants that have >90% VAF in T&NK cells were deemed to be germline variants and removed from the ratio calculation. The HSPC to T&NK variant ratio is calculated for each patient individually and different patient groups e.g. IM-sensitive, IM-resistant are compared. Additionally, mitochondrial variants were categorized as synonymous, non-synonymous, and non-coding, and the HSPC-to-T&NK variant ratio was calculated separately for each category. Finally, we generated UMAPs from the top 50 principal components (PCs) of the highly variable genes following the standard Seurat (v5.1) pipeline.

### Validation of mtDNA variant calling from 5**′** scRNA-seq data by comparison to targeted bulk sequencing

We validated mtDNA mutations identified in total white cells by confirming their presence in matched bulk CD34L leukemic stem/progenitor cells and by comparing variant calls from 5′ single-cell RNA sequencing data to targeted bulk mtDNA sequencing.^34,50^ Detailed methodology, variant calling performance metrics, and results are provided in the **Supplementary Methods** and **Supplementary Tables 7 and 8**.

### Statistical Analysis

Comparisons between groups were performed using Fisher’s exact test for categorical variables, and the Mann–Whitney U test or Kruskal–Wallis test for non-normally distributed continuous variables. For normally distributed data, ordinary one-way ANOVA was applied. Where applicable, Bonferroni correction was used to adjust for multiple comparisons. All statistical analyses were conducted using GraphPad Prism v9.0 (GraphPad Software, USA) and R v4.2.0 (R Foundation for Statistical Computing, Austria).

Kaplan–Meier estimates and log-rank tests were used to assess the probability of achieving MMR and MR4.5 at 24 months of TKI therapy, stratified by mtDNA mutational burden, D-loop and *MT-CO1* mutations.

Survival analyses, including overall survival (OS), progression-free survival (PFS), and failure-free survival (FFS), were estimated using the Kaplan–Meier method and compared using the log-rank test. FFS was defined according to ELN criteria, including failure to achieve time-dependent molecular milestones, acquisition of *BCR::ABL1* kinase domain mutations, progression to accelerated or blast phase, or death from any cause.

All p-values were two-sided, and values less than 0.05 were considered statistically significant.

## Acknowledgements

The authors thank the patients who kindly donated blood samples to make this research possible. We thank the staff of the South Australian Cancer Research Biobank, Princess Alexandra Hospital Cancer Collaborative Biobank and Australian Leukaemia and Lymphoma Group; and the South Australian Genomics Centre for the mtDNA targeted sequencing.

The authors are grateful to: Dr Monika Kutina and Dr Devendra Hiwase for the colony forming assay protocol.

This research project was supported by: SAHMRI Mid-Career Seed Funding Grant (I.S.P.); Cancer Council SA Research Mid-Career Research Fellowship on behalf of its donors and the State Government of South Australia through the Department of Health and Wellbeing (I.S.P.); Cancer Council SA Research COVID-19 Hardship Grant (I.S.P.); Investigator Grant # 2007908 (T.P.H); Contributing Haematologists Committee Research Grant (D.M.R, I.S.P., D.W). National Medical Research Council Singapore: CIRG/1468/2017, MOH-000602, MOH-000059, CIRG16nov032 (S.T.O.). Leukemia & Lymphoma Synergistic Team Award with support from the Mike & Sofia Segal Foundation (D.T.), NHMRC Ideas Grants (D.T.) and The Medical Research Future Fund (D.T.).

## Author Contributions

I.S.P. conceived the project, developed the concept, designed and performed experiments, analysed the data, and wrote the paper; V.K. performed the scRNA-seq experiments, analysed the sc data, generated figure 5, wrote the GSEA part of the paper and reviewed the manuscript; O.F.J. performed the bioinformatics analysis of the sc data, generated figure 5, wrote the sc analysis and reviewed the paper; C.H.K. performed the bioinformatics analysis of the targeted mtDNA mutations and reviewed the manuscript; B.C. performed the bioinformatics analysis of the sc data; K.L.L. performed the GSEA; C.M.P. and P.W. performed the validation analysis and generated Figure 2 C and D, and the pathogenicity score; E.P. performed the colony forming and MitoTracker experiments; P.D. assisted with the experiments; V.A.S. assisted with the experiments; J.J. performed the MitoTracker experiments; K.L. assisted with the Seahorse experiments; N.S. contributed clinical data and reviewed the manuscript; A.Y. contributed clinical data, reviewed the flow sorting gating and reviewed the manuscript; S.B. contributed clinical data and reviewed the manuscript; C.C. contributed clinical data and reviewed the manuscript; D.T.Y. contributed clinical data and reviewed the manuscript; D.W. contributed clinical data and reviewed the manuscript; V.G.H. shared protocols and reviewed the manuscript; D.T. shared protocols and reviewed the manuscript; T.P.H. reviewed the manuscript; S.T.O. reviewed the manuscript; M.M.R conceived the original project, designed experiments and wrote the paper.

## Data availability

We declare that data supporting the findings of this study are available within this manuscript and its supplementary information files.

Supplementary information accompanies the manuscript on the *Signal Transduction and Targeted Therapy* website http://www.nature.com/sigtrans

## Research Dataset

The 10X 5’ scRNA-seq data comprising CML CP patients with varying imatinib sensitivity is available as fastqc files at the EGA under accession ID: EGAS00001005509.

