## Supplemental Methods and Figures for "Mitochondrial DNA Mutations Determine Favourable Molecular Responses to Targeted Kinase Inhibitor Therapy and Impair Oxidative Phosphorylation"

**This PDF file includes:**

Materials and Methods
Figures. S1 to S5
Captions for Data S1 to S5

**Other Supplementary Materials for this manuscript include the following:**

Tables S1 to S8
S1: Characteristics of CML Patients at Study Entry
S2: GSEA of CD34+ HSPC
S3: Selected GSEA of CD34+ HSPC
S4: List of sc-mtDNA Variants
S5: mtDNA Somatic Mutations in Leukocytes
S6: mtDNA mutations in Seahorse patients
S7: Validated Somatic mtDNA Variants Detected in TWCs and CD34$^{+}$ Cells Using LoFreq and mtGATK
S8: Validation of sc-mtDNA variants

### Materials and Methods

#### Oxygen consumption rate (OCR) measurement

OCR was measured using the Seahorse XF96 Analyzer (Agilent Technologies, California, United States) with the Seahorse XF Cell Mito Stress Test Kit (Agilent, California, United States). Primary cells were cultured overnight in SFM with physiological growth factors and then resuspended in warm Seahorse XF RPMI medium (pH 7.4), supplemented with 10 mM glucose, 1 mM pyruvate, and 2 mM glutamine (all from Agilent Technologies, California, United States). Eight replicates were performed per sample.

A total of $1.2 \times 10^5$ cells (50 μL) were seeded per well of a Seahorse XF cell culture plate pre-coated with 22.4 μg/mL Corning® Cell-Tak™ Cell and Tissue Adhesive (Corning, New York, United States) and centrifuged at 800 rpm for 3 minutes without brakes. After centrifugation, warm complete Seahorse medium was gently added to each well, bringing the final volume to 180 μL. The cells were incubated at 37°C in a non-$CO_2$ incubator for 1 hour.

The cells were then transferred to the Seahorse analyzer, where OCR was measured at baseline and after subsequent injections of specific inhibitors. The first injection was of the ATP-synthase inhibitor oligomycin (1.5 μM), followed by the mitochondrial uncoupler carbonyl cyanide-4-(trifluoromethoxy)phenylhydrazone (FCCP, 2 μM), and rotenone and antimycin A (0.5 μM), inhibitors of complexes III and II, respectively. OCRs were normalized by cell number. These steps allowed for the measurement of the ATP-linked respiration after the injection of oligomycin, and calculated as the last rate measurement before oligomycin injection minus the minimum rate measurement after oligomycin injection, as well as maximal OCR (the highest rate of oxygen consumption achieved after injection of FCCP normalised on the minimum value after the injection of rotenone and antimycin A indicative of the non-mitochondrial oxygen consumption) and basal OCR (the rate of oxygen consumption under normal conditions, normalised on the non-mitochondrial oxygen consumption). The difference between maximal and basal OCR was used to calculate mitochondrial fitness, providing insights into mitochondrial function and capacity.

#### Mitochondrial content and membrane potential

Mitochondrial content and membrane potential were assessed using MitoTracker Green FM and MitoTracker Red CMXRos (Invitrogen *Carlsbad, California*) fluorescent probes, following the manufacturers' instructions.

Briefly, $1.2 \times 10^5$ cells were washed with cold 1× PBS and pelleted at 1200 rpm for 5 minutes. Cells were resuspended in pre-warmed (37 °C) 1× PBS supplemented with 2% FCS and transferred to a 96-well round-bottom plate. Cells were then incubated for 30 minutes at 37 °C in a humidified incubator with 5% $CO_2$, protected from light, with anti-human CD34–APC (clone 8G12, BD Biosciences, New Jersey, United States; 1:100), anti-human CD38–PE-Cy7 (clone HIT2, BD Biosciences; 1:100), MitoTracker Green FM (50 nM, Invitrogen *Carlsbad, California*), and MitoTracker Red CMXRos (5 nM, Invitrogen). DAPI (Sigma-Aldrich, Missouri, United States) was then added at a final concentration of 0.05 ug/mL, and cells were incubated for an additional 1 minute. Following staining, cells were washed with pre-warmed 1× PBS containing 2% FCS and pelleted at 1200 rpm for 5 minutes. The supernatant was discarded, and cells were resuspended in pre-warmed 1× PBS with 2% FCS. Fluorescence was acquired using a BD LSRFortessa X-20 Cell Analyzer, and data were analysed using FlowJo software v10.10.

**Next-generation sequencing of the entire mitochondrial genome**
MtDNA mutations were identified as previously described.[1] Briefly, genomic DNA was extracted using a phenol/chloroform method. The full 16.9 kb mitochondrial genome was amplified by long-range PCR using overlapping primer sets, and pooled amplicons were subjected to library preparation using the NexteraXT kit (Illumina, San Diego, CA)) and sequenced on an Illumina MiSeq platform.

The full 16.9 kb mtDNA was isolated by long-range PCR, generating two or three overlapping fragments. by using the primers: hmt F1 5'- AACCAAACCCCAAAGACACC-3', hmt R1 5'-GCCAATAATGACGTGAAGTCC-3' (amplicon length 9,289 bp), hmt F2 5'-TCCCACTCCTAAACACATCC-3', hmt R2 5'-TTTATGGGGTGATGTGAGCC-3' (7,626 bp); or Mito1 F 5'- ACATAGCACATTACAGTCAAATCCCTTCTCGTCCC-3', Mito1 R 5'-TGAGATTGTTTGGGCTACTGCTCGCAGTGC-3' (3,968 bp), Mito2 F 5'-TACTCAATCCTCTGATCAGGGTGAGCATCAAACTC-3', Mito2 R 5'-GCTTGGATTAAGGCGACAGCGATTTCTAGGATAGT-3' (5,513 bp), Mito3 F 5'-TCATTTTTATTGCCACAACTAACCTCCTCGGACTC-3', Mito3 R 5'-CGTGATGTCTTATTTAAGGGGAACGTGTGGGCTAT-3' (7,814 bp).[1,2]

The reaction mix contained 2.5 U Takara LA Taq (Clontech Laboratories, Mountain View, CA), 1X Takara LA PCR Buffer ll ($Mg^{2+}$plus), 0.2 mM dNTPs, 400 nM primers forward and reverse primers and 20 ng of human genomic DNA. PCR was performed using the following conditions: 95°C for two minutes; 30 cycles of 95°C for 15 sec and 68°C for 10 minutes; 68°C for 20 minutes. After PCR, the amplicons were quality checked on agarose gel electrophoresis, purified by either *ExoSAP*-IT (Affimetrix) or QIAquick Gel Extraction Kit (Qiagen), quantified by *Qubit® 3.0 Fluorometer* (Thermofisher), and equimolar pooled. Pooled amplicons (1 ng) were used for the library preparation by NexteraXT kit (Illumina, San Diego, CA) following the manufacturer instructions, and ran on Illumina MiSeq by using the Miseq Reagent v3 600 cycle kit.
The figure is generated with *Biorender.com*.

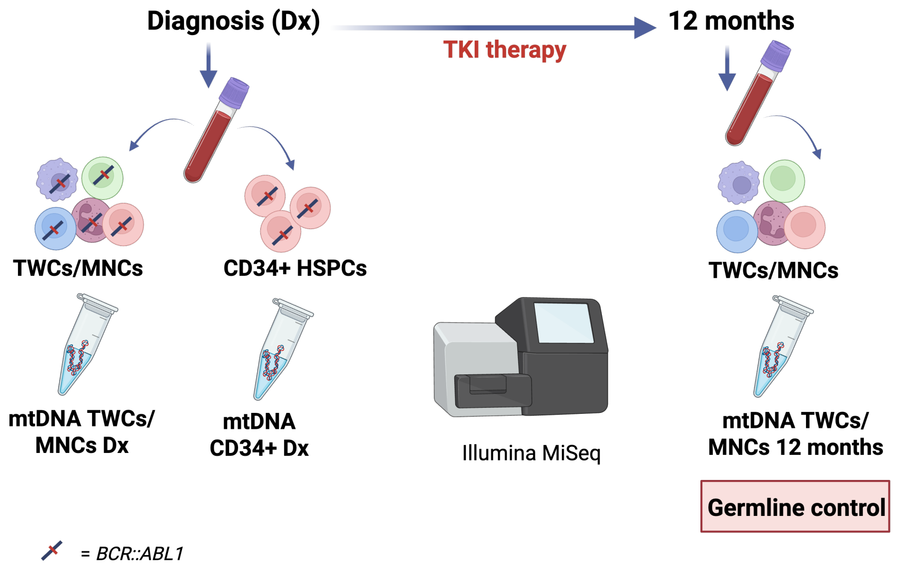

**MtDNA copy number quantification**

Relative mtDNA copy number was quantified by real-time quantitative PCR (qPCR) using two mitochondrial gene targets, MT-ND1 (Hs02596873-s1, FAM-MGB) and MT-CYB (Hs02596867-s1, FAM-MGB), normalized to the nuclear reference gene **GUS**B.[3] Reactions were performed in triplicate using TaqMan assays on a QuantStudio™ 7 Flex system (Life Technologies, California, USA).

All TaqMan® assays were obtained from Life Technologies (California, USA). Custom primers and probes for GUSB were: Forward: 5′-ATTTTGCCGATTTCATGACTGA-3′, Reverse: 5′-GACGGGTACGTTATCCCATGAG-3', Probe: FAM-5′-AGTGTAAGTGGCAGTTTG-3′-MGB.

Reactions were set up in 96-well Fast Optical Reaction plates containing 12.5 µL of TaqMan® Universal PCR Master Mix (Life Technologies), 400 nM of each primer, 200 nM of the GUSB MGB probe, 20 ng of genomic DNA, and nuclease-free water in a total volume of 25 µL. All reactions were performed in triplicate on a QuantStudio™ 7 Flex Real-Time PCR System (Applied Biosystems). Thermal cycling conditions were as follows: 2 min at 50 °C, 10 min at 95 °C, followed by 45 cycles of 15 s at 95 °C and 1 min at 60 °C. Ct values for MT-ND1, MT-CYB, and GUSB were determined simultaneously within the same run. Cases and matched controls were analysed on the same plate to minimize inter-assay variability. The relative mtDNA copy number per diploid genome was calculated using the formula: $MCN=2\times2^{-\Delta Ct}$, where $\Delta Ct$ = Ct_(MT-ND1 or MT-CYB) – Ct_(GUSB). The final MCN for each sample was determined by averaging the MCN values for both MT-ND1 and MT-CYB: Final $MCN=( MCN_{MT-ND1}+MCN_{MT-CYB})/2$.

**Validation of mtDNA variant calling from 5′ scRNA-seq data by comparison to targeted bulk sequencing**

We first confirmed that the mtDNA mutations identified in TWCs were also present in bulk CD34$^+$ LSPCs (Supplementary Table 7). Using LoFreq and mtGATK for variant calling, we validated six heteroplasmic somatic mutations initially detected in TWCs from four CP-CML patients at diagnosis but absent in matched germline controls (LoFreq: mean VAF 5.1%; mtGATK: mean 5.2%). These mutations were also identified in CD34$^+$ cells with comparable VAFs (LoFreq: mean 6.0%; mtGATK: mean 6.9%), supporting the persistence of clonally dominant mtDNA mutations in the LSPC compartment.

We next validated the mtDNA variant calling from 5′ scRNA-seq data by comparing them to those obtained from targeted mtDNA sequencing of matched bone marrow CD34$^+$ cells and mononuclear cells (MNCs) from the same patients using the Illumina MiSeq platform (**Supplementary table 8**). In bulk CD34$^+$ cells, a total of 29 homoplasmic mtDNA variants were identified using the LoFreq algorithm, while no heteroplasmic variants were detected. For scRNA-seq data, we applied a stringent cutoff of ≥2 cells and a minimum single-cell allele frequency (scAF) of 10% (scAF defined as variant-supporting reads divided by total mitochondrial reads per cell) to reduce false positives. Of the 29 validated variants, 13 were correctly identified (true positives), 16 were missed (false negatives), and 2 additional variants, with scAF of 91% and 96%, were detected that were not present in bulk data (false positives); 34 positions were correctly identified as negatives (true negatives). Although all mtDNA variants identified in CD34$^+$ bulk DNA were homoplasmic, their corresponding scAFs in scRNA-seq data showed a median of 96%

(range: 16–100%), with lower scAF associated with low coverage. This approach achieved a specificity of 94%, precision of 87%, and overall accuracy of 72%. However, sensitivity remained modest at 45%, with a false-negative rate of 55%, likely due to the intrinsic limitations of the 10x Genomics 5′ platform, including low mitochondrial coverage, read dropouts, and the transcript-based nature of the assay, which limits the ability to capture full-length mitochondrial genomes.[4,5]

**Figure. S1.**

**Mitochondrial function in CD34+ cells.**

Representative histograms of Mitotracker Green- and Mitotracker red CMRos-labelled normal CD34+ HSPCs (green) and CD34+ LSPCs (TKI-S in blue and TKI-Res in red) (Median Fluorescent Intensity).

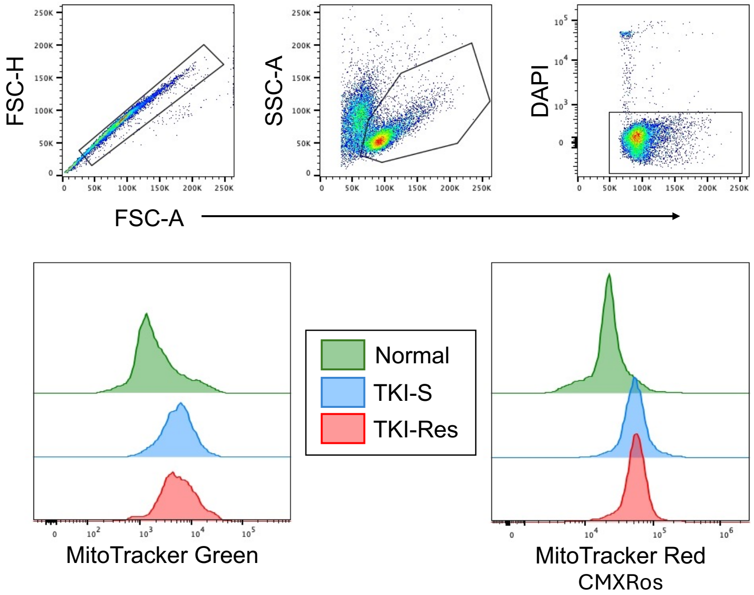

**Figure. S2.**

**Mitochondrial gene-set analyses in the 5' scRNA-seq dataset (ID: EGAS00001005509).**

**(A)** Mitochondrial gene-sets showing significant enrichment in gene set enrichment analysis (GSEA) (**Supplementary Table 6**) were subjected to leading-edge analysis. Genes identified in these pathways were compiled into a metagene list, and their expression was evaluated in the single-cell dataset using the "AddModuleScore" function. Module scores were pseudo-bulked per patient within the CD34+ LSPC compartment and visualized as box-and-whisker plots. Good responder: $CP^{MMR}$; Poor responders: $CP^{IMres}$ and $CP^{Fail}$. Statistical significance was assessed using a two-sided t-test ($p < 0.05$).

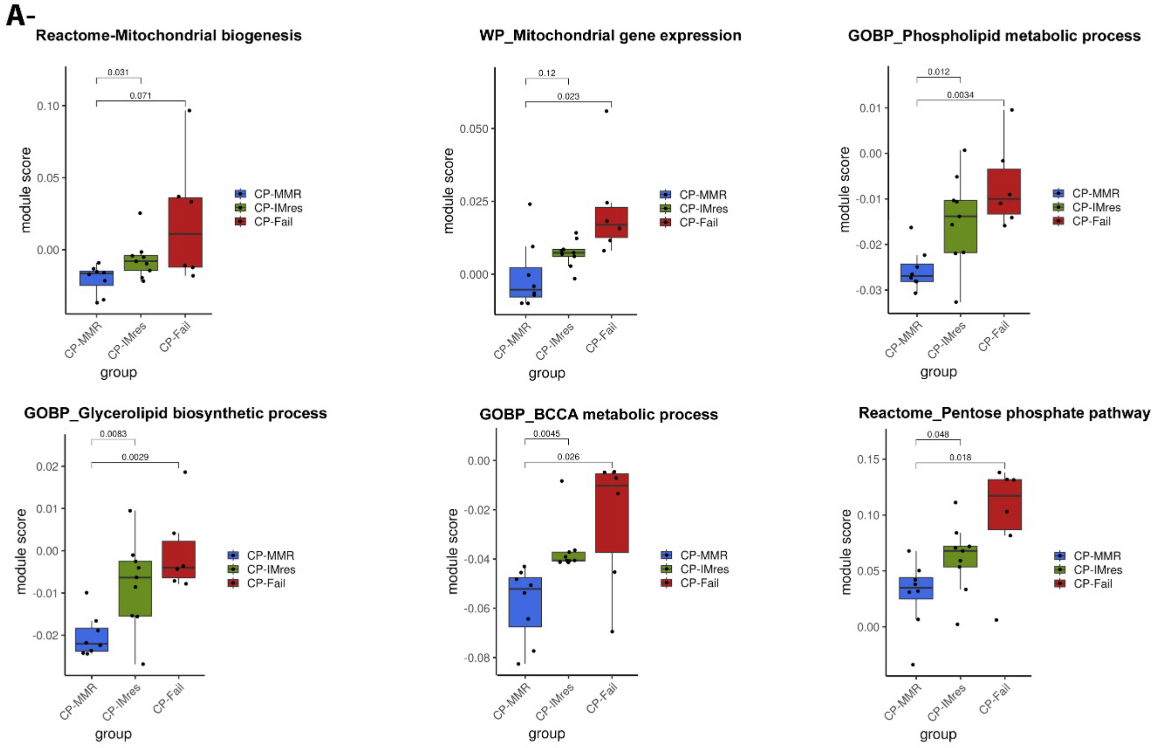

**Figure. S3.**

**Mitochondrial variant analysis in the 5' scRNA-seq dataset (ID: EGAS00001005509).**
**(A)** Coverage of scRNA-seq reads across mitochondrial genes for three representative samples: P147 (poor responder), P654, and P677 (good responders).

**(B-E)** Total mitochondrial variants detected in LSPC clusters (HSC, iMKP, MKP, ERP, EOBM, iNeP, NeP, Lyp, Pro-B) and T/NK cell clusters, stratified by variant type (**B**, all variants; **C**, synonymous; **D**, non-synonymous; **E**, non-coding). Data are shown per patient.

Abbreviations: *Hematopoietic stem cell (HSC), Immature megakaryocyte progenitor (iMKP), Megakaryocyte progenitor (MKP), Erythroid progenitor (ERP), Eosinophil-Basophil Mast cell progenitor (EOBM), Immature Neutrophil progenitor (iNeP), Neutrophil progenitor (NeP), Lymphoid progenitor (Lyp), Pro-B (B cell progenitor).*

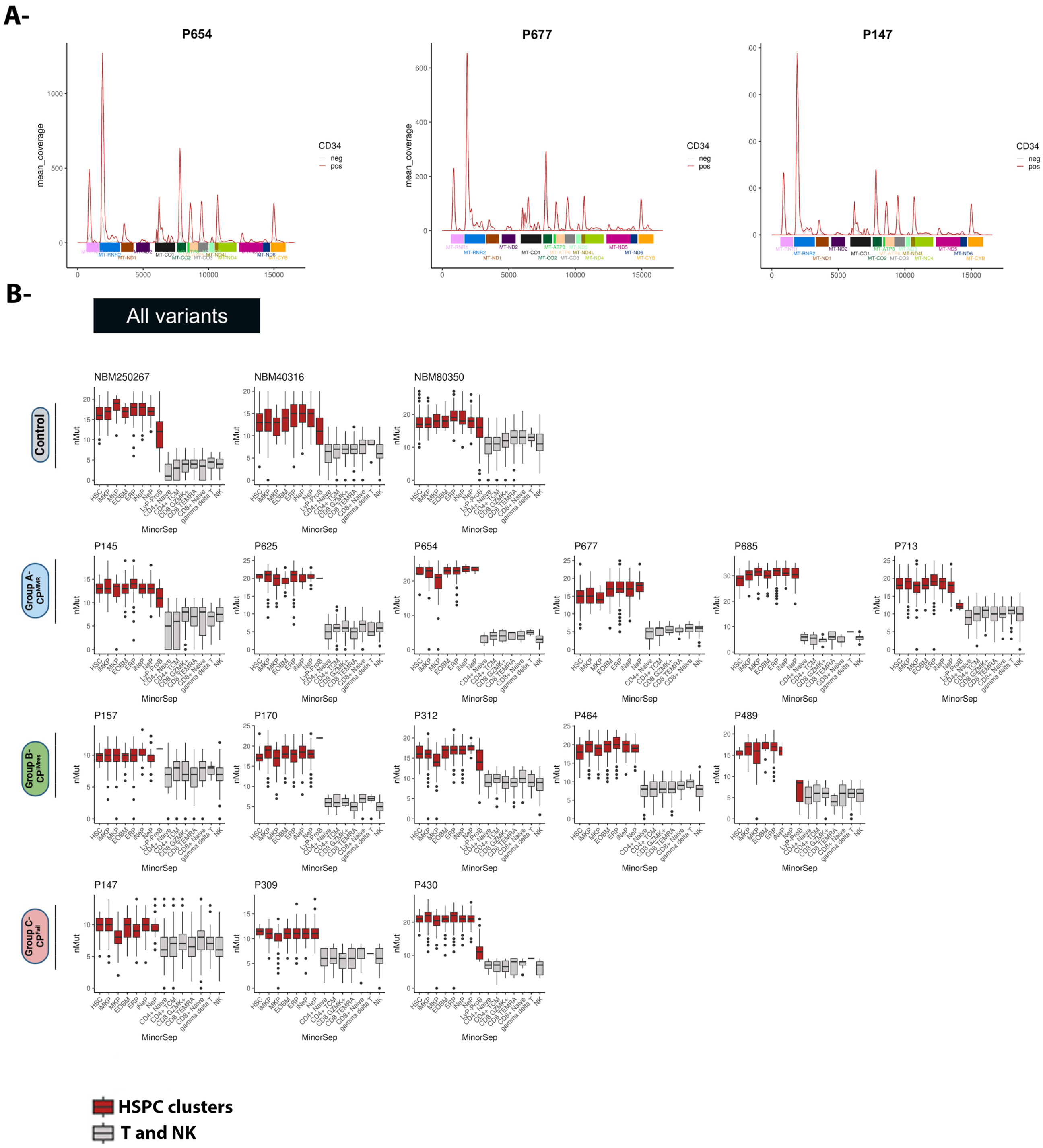
A-
P654
P677
P147
mean_coverage
CD34
neg
pos
B-
All variants
Control
NBM250267
NBM40316
NBM80350
nMut
MinorSep
P145
P625
P654
P677
P685
P713
P157
P170
P312
P464
P489
P147
P309
P430
HSPC clusters
T and NK

**C-**

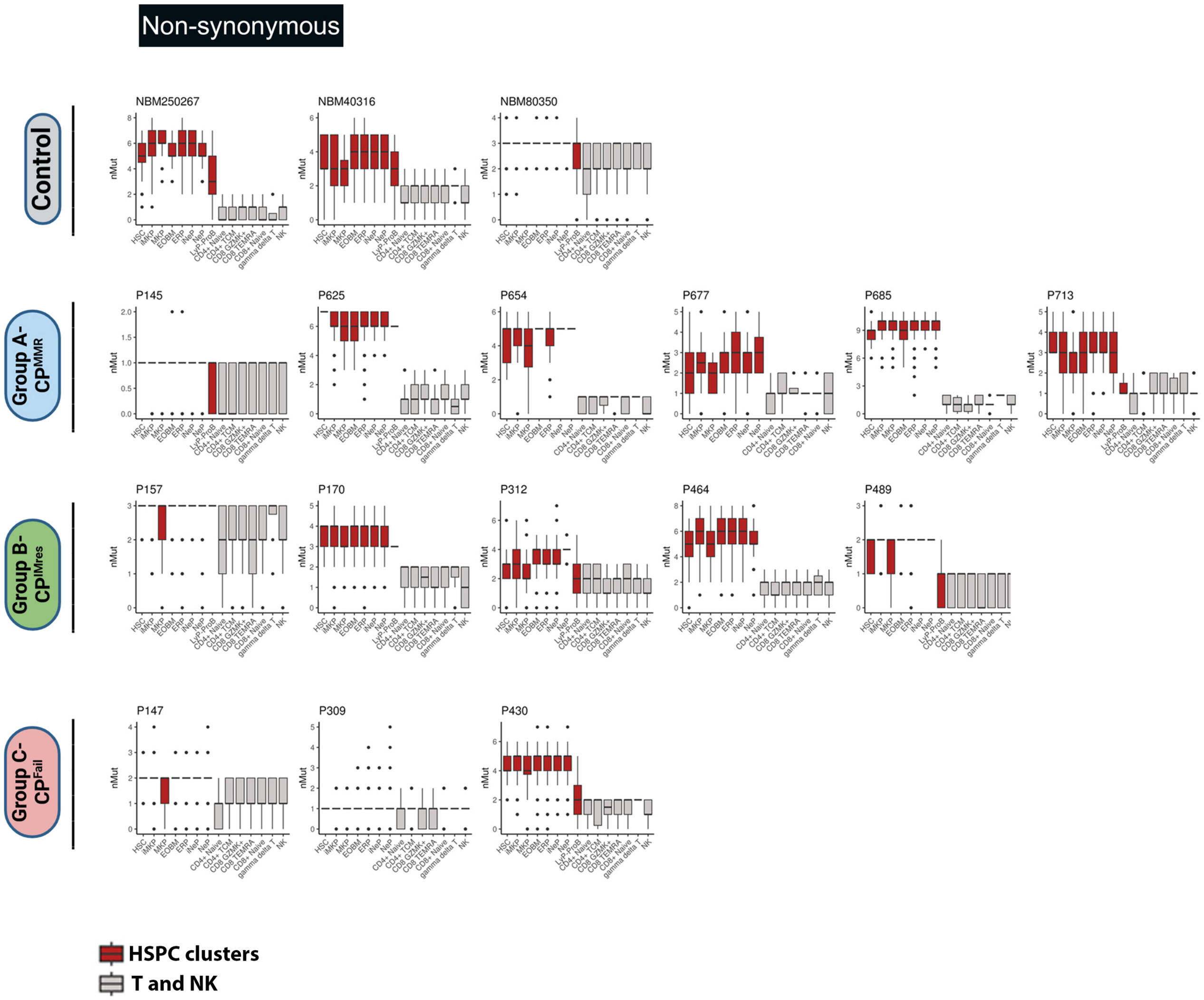
Non-synonymous
Control
NBM250267
NBM40316
NBM80350
Group A- CPMMR
P145
P625
P654
P677
P685
P713
Group B- CPIMres
P157
P170
P312
P464
P489
Group C- CPFail
P147
P309
P430
nMut
HSC
iMKP
MKP
EOBM
ERP
iNeP
NeP
LyP-ProB
CD4+ Naive
CD4+ TCM
CD8 GZMK+
CD8 TEMRA
CD8+ Naive
gamma delta T
NK
HSPC clusters
T and NK

D-

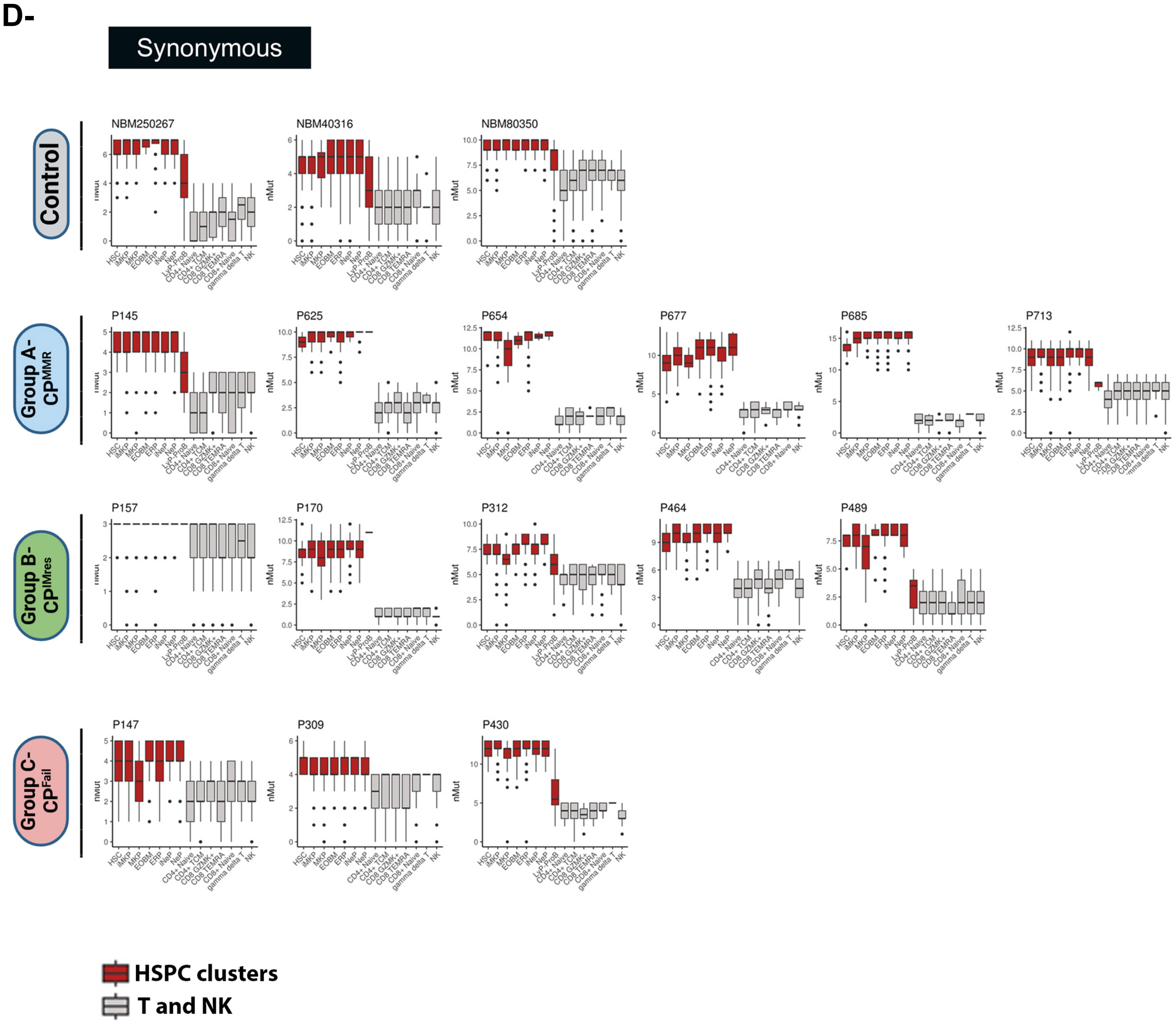

Synonymous
Control
Group A- CP^MMR
Group B- CP^IMres
Group C- CP^Fail
NBM250267
NBM40316
NBM80350
P145
P625
P654
P677
P685
P713
P157
P170
P312
P464
P489
P147
P309
P430
nMut
HSC
IMKP
MKP
EOBM
ERP
iNeP
NeP
LyP-ProB
CD4+ Naive
CD4+ TCM
CD8 GZMK+
CD8 TEMRA
CD8+ Naive
gamma delta T
NK
HSPC clusters
T and NK

**E-**

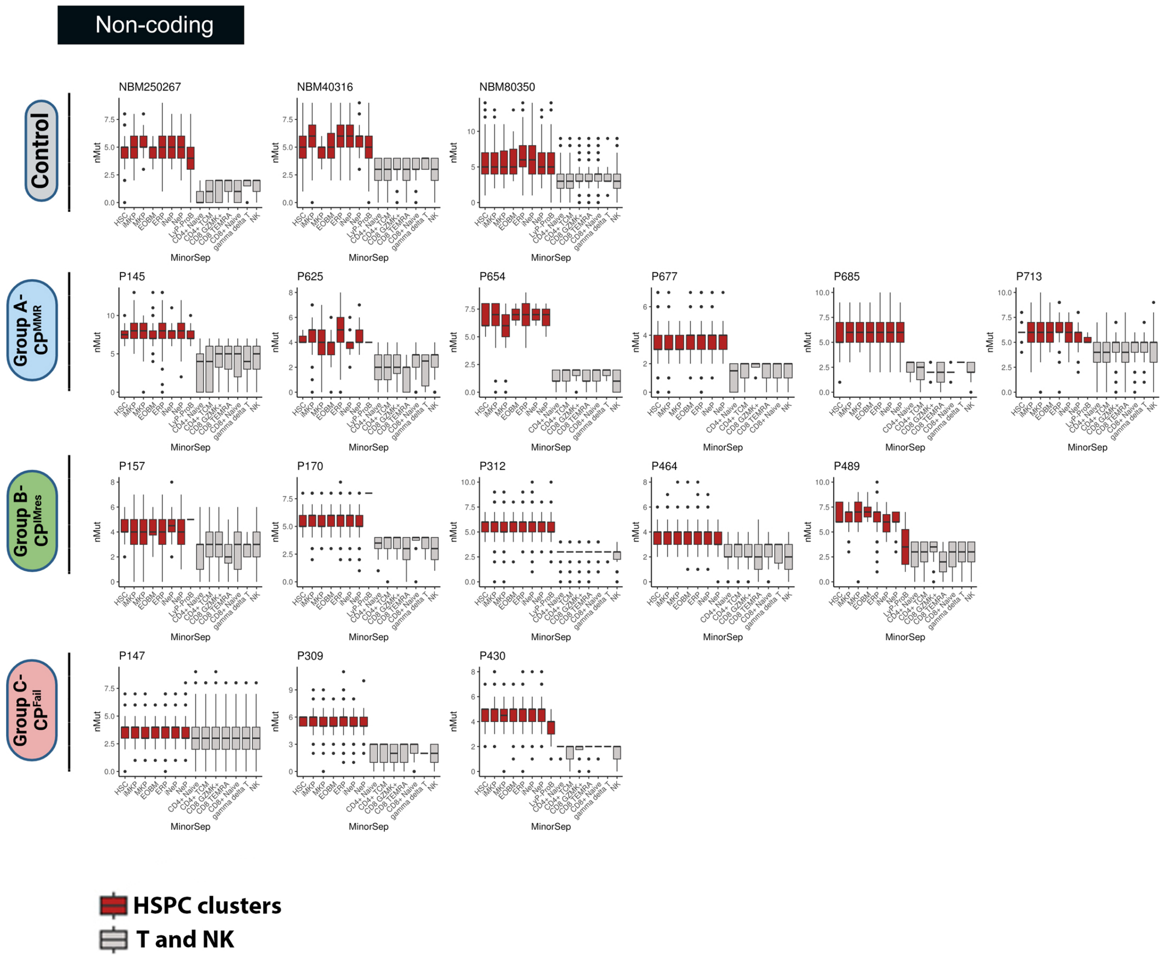
Non-coding
Control
Group A-CPMMR
Group B-CPIMres
Group C-CPFail
NBM250267
NBM40316
NBM80350
P145
P625
P654
P677
P685
P713
P157
P170
P312
P464
P489
P147
P309
P430
nMut
MinorSep
HSPC clusters
T and NK

**Figure. S4.**

**Association between mtDNA mutations and response to imatinib (IM)**

**(A)** Variant allele frequencies (VAFs) of structural RNA (r/tRNA) mutations in IM-sensitive (IM-S, blue) and IM-resistant (IM-Res, red) patients (Mann-Whitney test, median with interquartile range).

**(B)** Variant allele frequency (VAF) of synonymous mutations in IM-S vs. IM-Res patients (Mann-Whitney test, median with interquartile range).

**(C)** Distribution of PolyPhen-2 scores for non-synonymous mutations in IM-S and IM-Res patients (Mann-Whitney test, median with interquartile range). PolyPhen-2 scores >0.908 are classified as "probably damaging," scores between 0.446 and 0.908 as "possibly damaging," and scores <0.446 as "benign".

**(D)** Distribution of SIFT scores for non-synonymous mutations in IM-S and IM-Res patients (Mann-Whitney test, median with interquartile range). SIFT scores range from 0 to 1, with scores <0.05 considered deleterious and scores ≥0.05 considered tolerated.

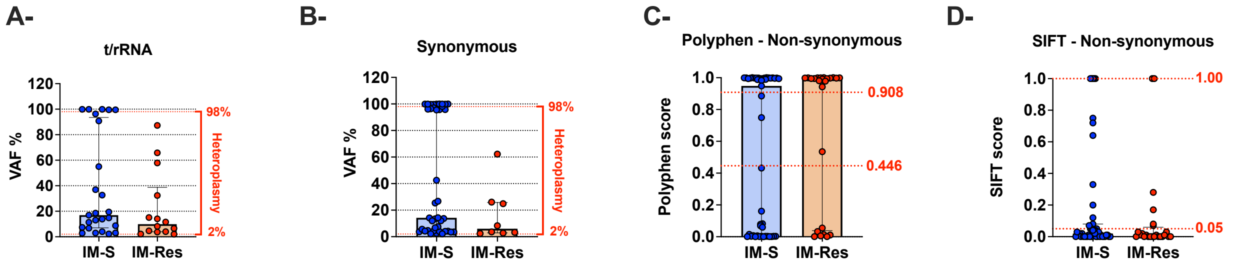

**Figure. S5.**

**Impact of mtDNA mutations on imatinib response.**

Failure-free survival (FFS) according to:

**(A)** D-Loop mutations vs non-D-loop at 2 years (100% vs 79%; hazard ratio log rank [HR] 0) and **(B)** at 4 years (100% vs 73%; hazard ratio log rank [HR] 0).

**(C)** mutations in MT-CO1 vs non-MT-CO1 mutations at 2 years (90% vs 80%; hazard ratio log rank [HR] 0.45; 95% CI, 0.09 – 2.03) and **(D)** at 4 years (90% vs 74%; hazard ratio log rank [HR] 0.39; 95% CI, 0.09 – 1.6).

**(E)** mutational burden, ≥3 or <3 mutations at 2 years (95% vs 75%; hazard ratio log rank [HR] 20.18; 95% CI, 0.05 – 0.59) and **(F)** at 4 years (82% vs 75%; hazard ratio log rank [HR] 0.54; 95% CI, 0.18 – 1.63).

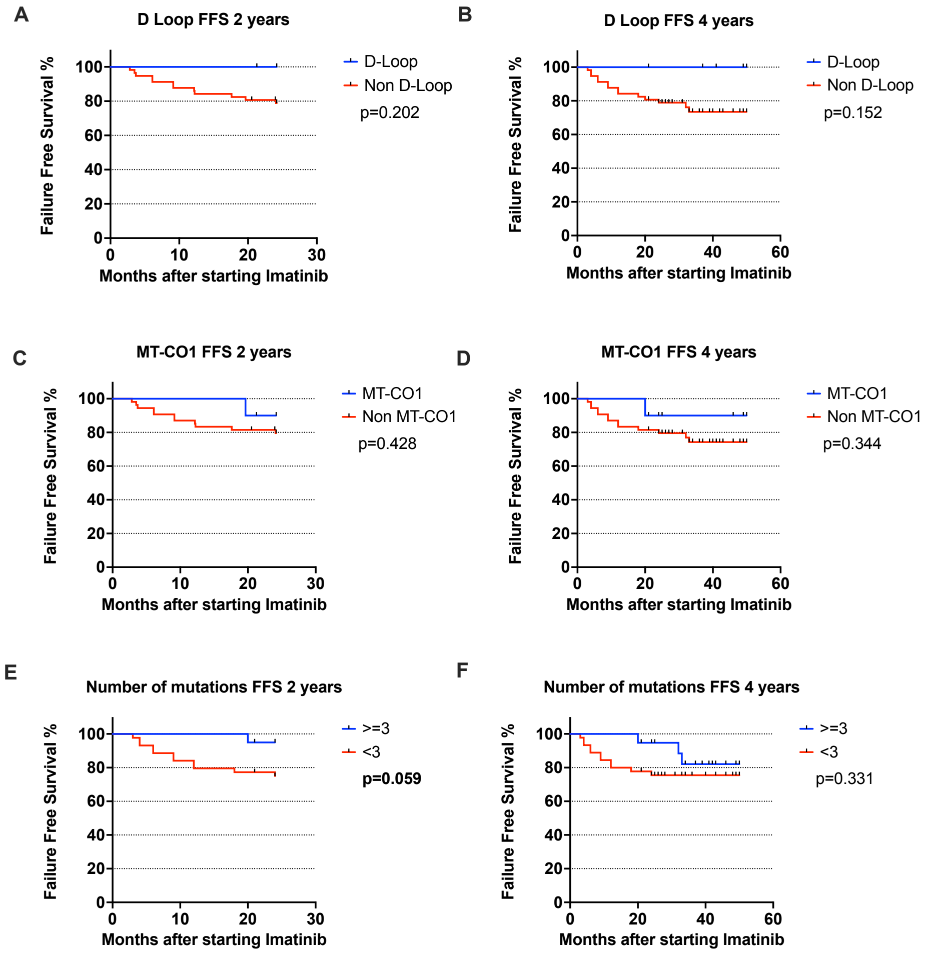
